# Filopodia-mediated trans-endocytosis

**DOI:** 10.64898/2026.02.09.703982

**Authors:** Hanna Grobe, Marcela Xiomara Rivera Pineda, Sujan Ghimire, Marie-Catherine Laisne, Anna Nylund, Monika Vaitkevičiūtė, Helena Vihinen, Anu Prakash, Johanna Tammi, Marjaana Ojalill, Pia Boström, Pauliina Hartiala, Johanna Englund, Emilia Peuhu, Eija Jokitalo, Guillaume Jacquemet

## Abstract

Cell–cell communication in tissues is influenced by contact geometry and molecular signalling. Here, we demonstrate that epithelial intercellular filopodia can penetrate neighbouring cells to form micrometre-long, double-membrane “impact sites” and, in some instances, initiate trans-endocytosis. Using super-resolution live imaging and three-dimensional electron microscopy in cell models, xenografts, and patient-derived tissues, we observe widespread intercellular filopodia and extensive interdigitation at the cell-cell interface. Filopodia impact sites in the recipient cell recruit PACSIN2 into dynamic “PACSIN2 fingers” and are specifically enriched for caveolin-1 and dynamin-2, while clathrin and CLIC pathway markers are not enriched. Most contacts are transient and resolved by retraction, but some undergo scission, with recipient cells internalising filopodial tips across both homotypic and heterotypic interactions, including cancer–endothelium contacts. Correlative light and FIB–SEM analysis reveals that internalised tips are often double-membraned, closely associated with endoplasmic reticulum tubules, and can traffic to lysosomes. These findings define a protrusion-driven, curvature-dependent uptake pathway at cell–cell interfaces and identify filopodia as exchange organelles in epithelial collectives.

## Introduction

In multicellular organisms, cells exist within densely inter-connected communities in which coordinated tissue-level behaviours emerge from communication with both the extracellular matrix (ECM) and neighbouring cells. In particular, cell-cell signalling operates across various spatial scales, with diffusible cues such as chemokines and extracellular vesicles enabling long-range communications (Haas and Gilmour, 2006; Hoshino et al., 2015). At shorter distances, contact-dependent signalling can occur through juxtacrine receptor–ligand interactions and cell–cell adhesion receptors. Cells also communicate mechanically (Biggs et al., 2020), with adhesion complexes and the cytoskeleton transmitting forces over multiple cell lengths, thus enabling coordinated behaviours such as morphogenesis, repair, and collective migration (Sunyer et al., 2016). These biochemical and mechanical networks are closely integrated, and their dysregulation can amplify pathological signalling and disrupt tissue architecture in diseases, including cancer (Su et al., 2024).

Contact-dependent cell–cell communication depends not only on molecular identity but also on geometry, including which cells can be reached, where contacts form, and how long they last. Filopodia, thin actin-rich projections, extend a cell’s physical reach. They provide extended contact sites where adhesion and signalling receptors gather, supporting signal exchange and the initiation or reinforcement of adhesions (Ruhoff et al., 2025). For example, in epithelial cells, Ca^2+^-induced filopodia promote E-cadherin clustering into “zipper”-like adhesive arrays and the formation of adherens junctions (Vasioukhin et al., 2000). During embryogenesis, E-cadherin–dependent filopodia drive cell-shape changes necessary for embryo compaction (Fierro-González et al., 2013). During wound healing and morphogenetic closure, filopodia at the advancing epithelial fronts can mediate “zippering” of opposing sheets and ensure proper tissue alignment (Abreu-Blanco et al., 2012; Jacinto et al., 2000; Millard and Martin, 2008; Zhang et al., 2018). Beyond these short-range filopodia adhesion-inducing roles, specialised filopodia-like protrusions, called cytonemes, transport proteins and facilitate the exchange of membrane components across micrometre distances, thereby creating direct long communication pathways between cells (Brunt et al., 2021; Sanders et al., 2013). Collectively, these observations establish filopodia as active organisers of contact-dependent cell-cell communications and signalling.

In cancer, filopodia are often upregulated and associated with invasive behaviour (Jacquemet et al., 2015). However, most research has centred on filopodia–ECM interactions, where filopodia explore cell substrate topography and stiffness (Ruhoff et al., 2025), form adhesions (Jacquemet et al., 2016, 2019), remodel the extracellular matrix (Corinus et al., 2026; Peuhu et al., 2022; Summerbell et al., 2020), and direct migration and invasion (Arjonen et al., 2014; Li et al., 2014). In contrast, the development, organisation and fate of filopodia that engage neighbouring cells, especially within dense, dynamic collectives where membranes are closely opposed and constantly remodelled, remain largely unstudied.

Here, we examined the role of filopodia at the basolateral cell–cell interfaces in pre-invasive breast cancer, ductal car-cinoma in situ (DCIS), where tumour cells remain closely packed and confined within the basement membrane. We observe widespread intercellular filopodia in vitro, in DCIS-like in vivo xenografts, and in patient-derived DCIS tumour tissue. Using high-resolution live imaging and three-dimensional electron microscopy, we show that these intercellular filopodia can penetrate neighbouring cells, causing notable membrane invaginations that we term filopodia impact sites. We find that the F-BAR protein PACSIN2 (Protein Kinase C and Casein Kinase Substrate in Neurons Protein 2), along with CAV1 (caveolin-1) and DNM2 (dynamin-2), are recruited to these impact sites. We also report that, while most filopodia retract, some filopodia undergo trans-endocytosis, allowing direct engulfment of material by the adjacent cell. Internalised filopodial material then associates with endoplasmic reticulum tubules and can be trafficked to lysosomes for degradation. Overall, our data provide evidence for trans-endocytosis triggered by filopodial structures, revealing a previously underappreciated route for contact-dependent cell-cell communication and material exchange that may be relevant to tissue organisation and cancer progression.

## Results

### Intercellular filopodia are a conserved characteristic of epithelial cell–cell interfaces

We previously reported prominent filopodia at the leading edge of collectively migrating DCIS.com cells, a cellular model of DCIS, where they contribute to invasive migration (Peuhu et al., 2022; Jacquemet et al., 2017). In addition to these leading-edge structures, we observed abundant filopodia-like protrusions forming in between neighboring cells within migrating monolayers (Fig. 1A). Structured illumination microscopy (SIM) confirmed that these intercellular filopodia are present at basal, lateral, and apical surfaces along cell-cell interfaces (Fig. 1B). High-resolution live cell imaging further revealed that basolateral intercellular filopodia are highly dynamic, undergoing rapid cycles of protrusion, bending, and retraction during collective migration (Fig. 1C; Video 1). Consistent with these observations, live Airyscan imaging showed that intercellular filopodia repeatedly extend into and occupy transient gaps between cells, continuously remodelling as the monolayer migrates (Fig. S1A; Video 1). Ultrastructural analysis by electron microscopy (EM) confirmed the presence of intercellular filopodia within DCIS.com cell monolayers and further indicated that these protrusions can induce local membrane deformations in neighbouring cells (Fig. 1D; Fig. S1B).

**Fig. 1.**
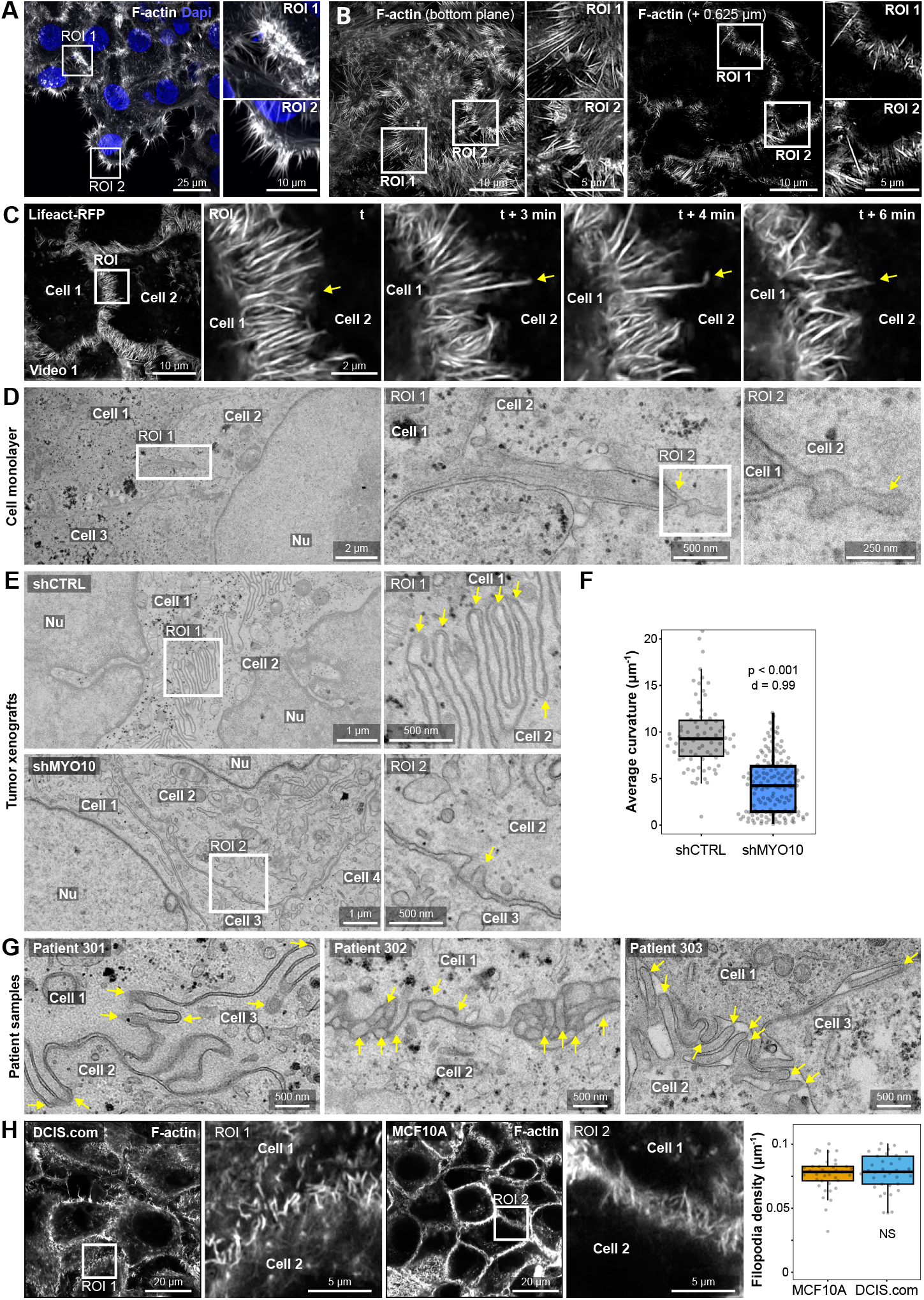
Intercellular filopodia are observed in vitro, DCIS-like xenografts, and patient-derived DCIS lesions. (**A**) DCIS.com cells were seeded in circular invasion assays and allowed to invade into fibrillar collagen I for 24 hours. Cells were fixed, stained with phalloidin (F-actin) and DAPI (nuclei), and imaged with spinning disk confocal microscopy (100× objective, CMOS camera). Representative maximum-intensity Z-projection are displayed. (**B**) DCIS.com cells were prepared as in (**A**), fixed, stained for F-actin, and imaged using structured illumination microscopy. Representative single Z planes highlighting filopodia at basal and lateral cell surfaces, with magnified ROIs, are presented. (**C**) DCIS.com lifeact-RFP cells were prepared as in (**A**) and imaged live with SIM. Video quality was enhanced using CSBDeep CARE (see methods). A yellow arrow indicates a representative filopodium undergoing protrusion, bending, and retraction (see **Video 1**). (**D**) DCIS.com cell monolayers were fixed and examined with electron microscopy (EM). A representative EM image depicts intercellular filopodia and the membrane deformation they induce, with magnified views provided. (**E, F**) shCTRL and shMYO10 DCIS-like xenografts were imaged 25 days post-inoculation with EM to visualise the cell-cell interface within the acini. (**E**) Representative images and magnified ROIs are shown. (**F**) Cell-cell interface curvature was quantified using the Fiji plugin “Kappa” based on manual tracing of EM images (n > 79 fields of view, 3 biological repeats, >22 fields per repeat). (**G**) Patient-derived breast tissues processed and imaged by EM display representative examples of intercellular filopodia at cell-cell interfaces in each patient. (**H**) DCIS.com and MCF10A cell monolayers were fixed, stained for F-actin, and imaged using Airyscan confocal microscopy. Representative single Z-plane images are shown. Filopodia density at cell-cell contact was quantified (see methods) (n > 37 FOV, 3 biological repeats). (**F, H**) Results are shown as boxplots, with whiskers extending from the 10th to the 90th percentiles. The boxes indicate the interquartile range, and a line within each box marks the median. Data points outside the whiskers are shown as individual dots. The raw numerical values and images used to make this figure have been archived on Zenodo (Grobe et al., 2026).

To determine whether these structures are also present in vivo, we analysed DCIS-like tumour xenografts generated from control DCIS.com cells (shCTRL) or from cells with downregulated Myosin-X (MYO10) level leading to reduced filopodia formation (shMYO10) (Peuhu et al., 2022). EM imaging showed complex, highly interdigitated cell-cell interfaces within shCTRL xenografts at both early (10-day) and late (25-day) stages, with prominent filopodial protrusions already visible at day 10 (Fig. S1C) and significantly increased interface complexity at day 25 (Fig. 1E; Fig. S1D). Conversely, shMYO10 xenografts displayed visibly reduced interdigitation, indicated by a notable decrease in cell-cell interface curvature (Fig. 1E, F). These findings support the conclusion that intercellular filopodia are present in vivo and that MYO10-dependent filopodia contribute to the formation of complex interdigitated interfaces within DCIS-like lesions.

We next examined patient-derived breast tissues collected from three individuals undergoing mastectomy for DCIS. EM analysis revealed intercellular filopodia and membrane inter-digitations at cell-cell interfaces across all samples (Fig. 1G; Fig. S2A). Notably, the sample from patient 303, whose diagnostic samples ultimately contained no traces of tumour despite the earlier suspicion of DCIS, exhibited markedly simpler cell-cell interfaces and reduced curvature compared to the confirmed DCIS samples 301 and 302 (Fig. S2B). Although the limited cohort precludes firm conclusions regarding clinical relevance, these observations prompted us to investigate them further in vitro using non-transformed (MCF10A) and transformed (DCIS.com) epithelial cells. Airyscan imaging revealed comparable densities of intercellular filopodia at basolateral cell-cell contacts in both cell lines (Fig. 1H), indicating that intercellular filopodia formation is not restricted to transformed cells. Finally, EM analysis of spheroids generated from MCF10A and T-47D cells using two independent approaches (Cultrex embedding or low-attachment culture) revealed filopodia-like protrusions at cell-cell interfaces across all conditions (Fig. S3), demonstrating that intercellular filopodia are also a feature of three-dimensional (3D) epithelial organisation.

In summary, intercellular filopodia represent a conserved and prominent feature of epithelial cell–cell interfaces in 2D migration assays, 3D spheroids, DCIS-like xenografts, and patient-derived breast tissues.

### Filopodia can penetrate and deform membranes at intercellular impact sites

The complex architecture of cell-cell interfaces (Fig. 1) prompted us to examine these regions at ultrastructural resolution using focused ion beam scanning electron microscopy (FIB-SEM; Fig. 2A).

**Fig. 2.**
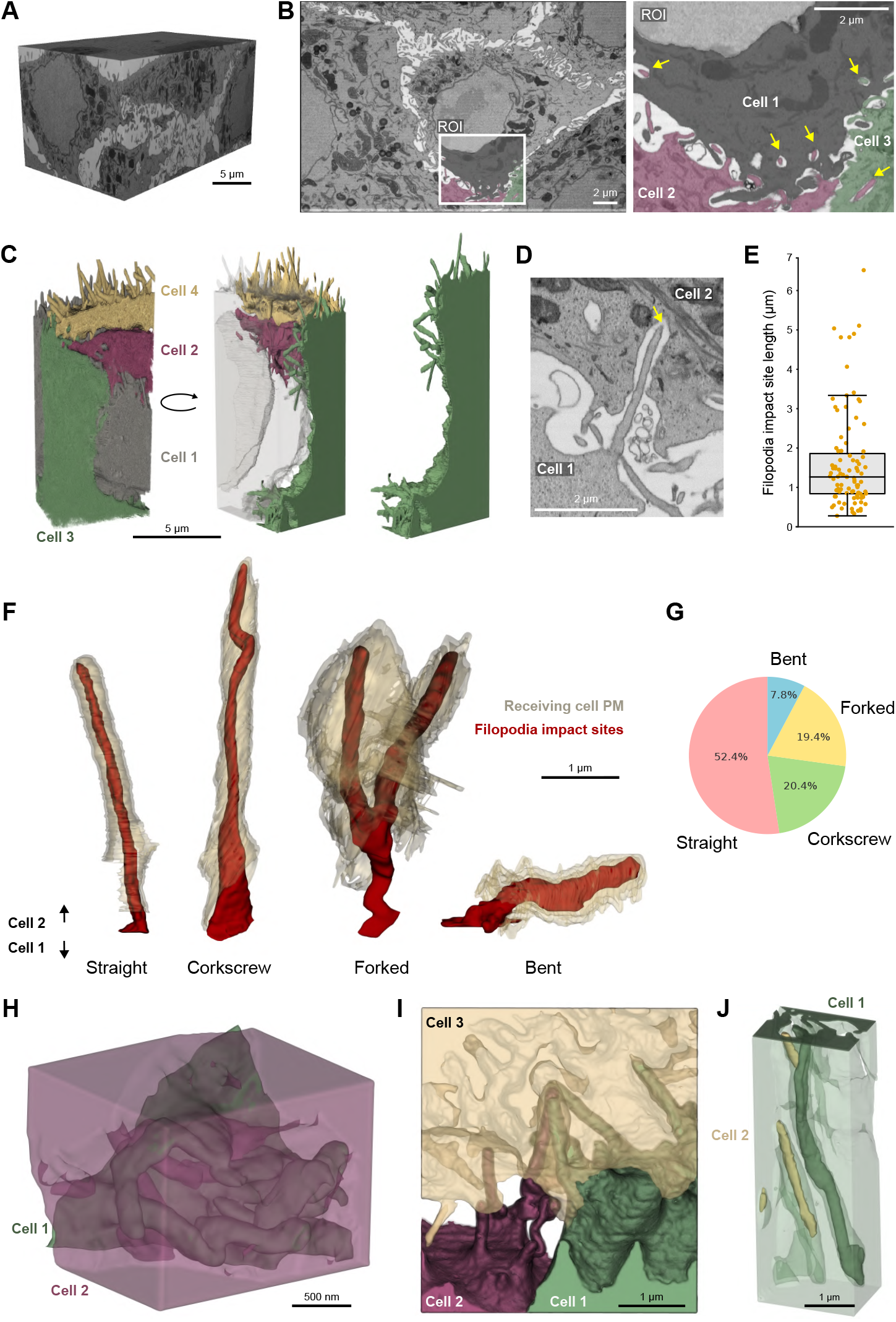
FIB-SEM reveals filopodia penetration and impact sites at the DCIS.com cell-cell interface. (**A-J**) A DCIS.com monolayer was fixed and processed for focused-ion-beam scanning electron microscopy (FIB-SEM). (**A**) Volume rendering of one representative FIB-SEM dataset. (**B, C**) To characterise the DCIS.com cell-cell interface in detail, a subvolume of the dataset was fully manually segmented. (**B**) The segmented region is highlighted. (**C**) Three-dimensional reconstruction of the segmented structures, allowing assignment of the cellular origin of individual filopodia at the cell-cell interface. See also **Video 2-4**. (**D**) Representative two-dimensional FIB-SEM slice showing a filopodium penetrating a neighbouring cell and forming a filopodia impact site. (**E**) Quantification of the length of filopodia impact sites (2 FIB-SEM volumes from 2 independent experiments, n = 100 filopodia). Results are shown as boxplots, with whiskers extending from the 10th to the 90th percentiles. The boxes indicate the interquartile range, and a line within each box marks the median. Data points outside the whiskers are shown as individual dots. (**F, G**) Representative segmentations of filopodia impact sites exhibiting distinct morphologies, including straight, corkscrew, forked, and bent structures (**F**), and their relative frequency of occurrence is shown as a pie chart (**G**) (2 FIB-SEM volume from 2 independent experiments, n = 100 filopodia). (**H-J**) Additional structural features observed upon detailed inspection of the FIB-SEM dataset, including the presence of “filopodia caves” (**H**), impact sites formed by filopodia originating from multiple cells (**I**), and cis impact sites in which filopodia fold back into the cell from which they originate (**J**). See also **Video 4**. The numerical data and raw images used to generate this figure have been archived on Zenodo (Grobe et al., 2026).

Three-dimensional FIB-SEM imaging of DCIS.com mono-layers revealed extensive filopodia-mediated interactions at cell-cell contact sites. To identify the cellular origin and organization of individual filopodia at these interfaces, we manually segmented a representative subvolume of the dataset (Fig. 2B), enabling three-dimensional reconstruction of cell boundaries and determination of filopodial origins (Fig. 2C). This analysis showed that filopodia originating from one cell often penetrate neighbouring cells, forming well-defined filopodia impact sites associated with notable double-membrane deformations (Fig. 2C, D). Quantitative analysis indicated that most impact sites extend 1–2 µm into receiving cells, with some reaching up to 6 µm (Fig. 2E).

Higher-magnification inspection revealed frequent membrane bulges in recipient cells at filopodia-impact sites (Fig. S4A). Quantification across independent FIB-SEM volumes showed that individual impact sites often contain multiple membrane bulbs (Fig. S4B), emphasising the structural complexity of these interfaces.

Morphological classification of filopodia impact sites revealed significant heterogeneity, with filopodia adopting straight, bent, corkscrew-like, or forked configurations (Fig. 2F). Straight filopodia were the most common (Fig. 2G). Corkscrew-like configurations resemble filopodia buckling observed in other contexts (Leijnse et al., 2015, 2022). In addition to these typical structures, we identified several less frequent but notable features, including cave-like membrane invaginations associated with multiple filopodial penetrations (filopodia caves; Fig. 2H), impact sites shared by filopodia originating from multiple neighbouring cells (Fig. 2I), and cis impact sites where filopodia fold back and invaginate into their originating cell (Fig. 2J).

These findings demonstrate that filopodia-like protrusions can induce deep, extensive membrane deformations at intercellular contact sites, revealing a previously unappreciated structural complexity at filopodia-mediated cell–cell interfaces.

### The F-bar protein PACSIN2 is recruited to filopodia impact sites

Filopodia impact sites exhibit striking membrane curvature on the recipient cell, suggesting that curvature-sensing scaffolds may be recruited to these sites. A previous study indicated that the F-BAR protein PACSIN2 (Protein Kinase C and Casein Kinase Substrate in Neurons Protein 2) gathers at membrane invagination sites during the intercellular spread of Listeria (Sanderlin et al., 2019). Given the structural similarities between Listeria-induced actin tail invasion and filopodia-driven membrane deformation, we hypothesised that PACSIN2 might also localise to filopodia impact sites.

To investigate this, we transfected DCIS.com lifeact-RFP cells with PACSIN2-GFP and conducted live cell imaging using Airyscan confocal microscopy. We observed dynamic, PACSIN2-enriched structures, hereafter referred to as “PACSIN2 fingers,” that specifically formed at filopodia impact sites along cell-cell interfaces (Fig. 3A and Video 5). To conclusively determine the origin of these structures, we co-cultured DCIS.com parental cells transiently expressing PACSIN2-GFP with DCIS.com lifeact-RFP cells. PACSIN2 consistently accumulated at contact points in recipient cells where filopodia from lifeact-RFP-positive neighbours extended (Fig. S5A and Video 5).

**Fig. 3.**
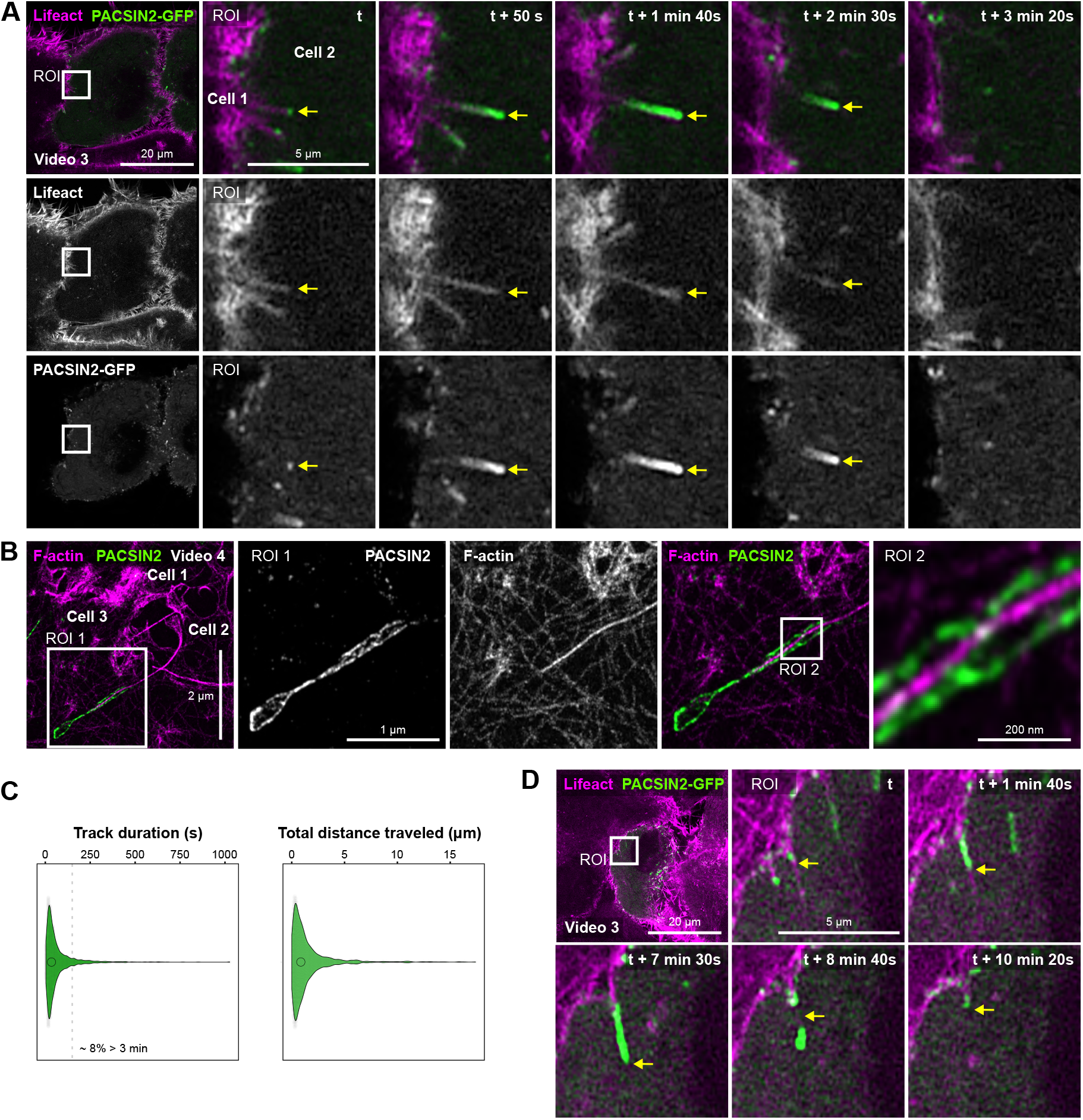
PACSIN2 is recruited to filopodia impact sites. (**A**) DCIS.com lifeact-RFP cells transiently expressing PACSIN2-GFP were cultured to form a monolayer and imaged live using Airyscan confocal microscopy. While all cells express lifeact-RFP to label F-actin, only a subset of cells express PACSIN2-GFP. A representative field of view and a magnified region of interest (ROI) are shown. The yellow arrow marks the recruitment of PACSIN2 to a filopodia impact site. (**B**) Monolayers of DCIS.com cells were fixed, stained for endogenous PACSIN2 and F-actin, and processed for 10x expansion microscopy. The expanded samples were imaged using Airyscan confocal microscopy. A representative single Z-plane and magnified ROIs show PACSIN2 localising around an actin-rich filopodium. (**C**) Dynamics of PACSIN2-positive structures (“PACSIN2 fingers”) were tracked from live imaging experiments. Violin plots display the distribution of track durations and total distances travelled (n = 1982 structures from 6 videos). (**D**) Live imaging of DCIS.com lifeact-RFP cells transiently expressing PACSIN2-GFP shows a long-lived PACSIN2-positive filopodium. A representative field of view and magnified ROI are shown. The yellow arrow highlights a stable PACSIN2-enriched structure that eventually breaks and is internalised over the course of the video. The raw images used to make this figure have been archived on Zenodo (Grobe et al., 2026).

Next, we examined whether endogenous PACSIN2 was similarly recruited to filopodia impact sites. Immunofluorescence staining of non-transfected monolayers followed by Airyscan imaging confirmed colocalisation of endogenous PACSIN2 with F-actin-positive filopodia at basolateral cell-cell interfaces (Fig. S5B), supporting the physiological relevance of our findings. As seen with electron microscopy, these PACSIN2 fingers can be long (Fig. S5C). To achieve higher resolution and better visualise PACSIN2 localisation, we used 10x expansion microscopy combined with Airyscan imaging (Damstra et al., 2022). This approach revealed PACSIN2 structures surrounding the penetrating filopodia without direct overlap (Fig. 3B and Video 6). Interestingly, the PACSIN2-positive area extended beyond the length of the actin-rich filopodia, ending in a distinct bud-like structure (Fig. 3B and Video 6), similar to what was observed with electron microscopy (Fig. 1D).

To understand the temporal behaviour of PACSIN2-positive fingers, we tracked PACSIN2-positive structures over time. Most PACSIN2 fingers were short-lived, with a median life-time of approximately 30 seconds, although about 8% persisted for longer than 3 minutes. Some even lasted up to 15 minutes (Fig. 3C). Remarkably, live imaging showed that a subset of these longer-lasting structures eventually disassembled and were internalised into the recipient cell (Fig. 3D and Video 5), likely through trans-endocytic membrane uptake.

### Cells can internalise intercellular filopodia

To investigate the fate of intercellular filopodia, we created a DCIS.com cell line that stably expresses fluorescently labelled filopodia tips. Since full-length MYO10-GFP exceeds the lentiviral packaging capacity and can artificially induce filopodia formation, we instead expressed a truncated MYO10 construct containing only the head, neck, and coiled-coil domains, called MYO10^HMM^-GFP (Fig. S6A) (Berg and Cheney, 2002). In DCIS.com cells, MYO10^HMM^-GFP was strongly localised to the tips of filopodia (Fig. S6B), and did not affect filopodia density or cell spreading compared to control cells (Fig. S6C and S6D).

Next, we assessed the localisation of MYO10^HMM^-GFP at cell-cell interfaces using Airyscan microscopy. Imaging revealed clear enrichment of MYO10^HMM^-GFP at the distal ends of PACSIN2-positive structures, confirming their filopodial identity (Fig. 4A, B and Fig. S7A). In these experiments, we also observed intracellular MYO10^HMM^-GFP puncta (Fig. 4A), prompting us to further investigate their origin.

**Fig. 4.**
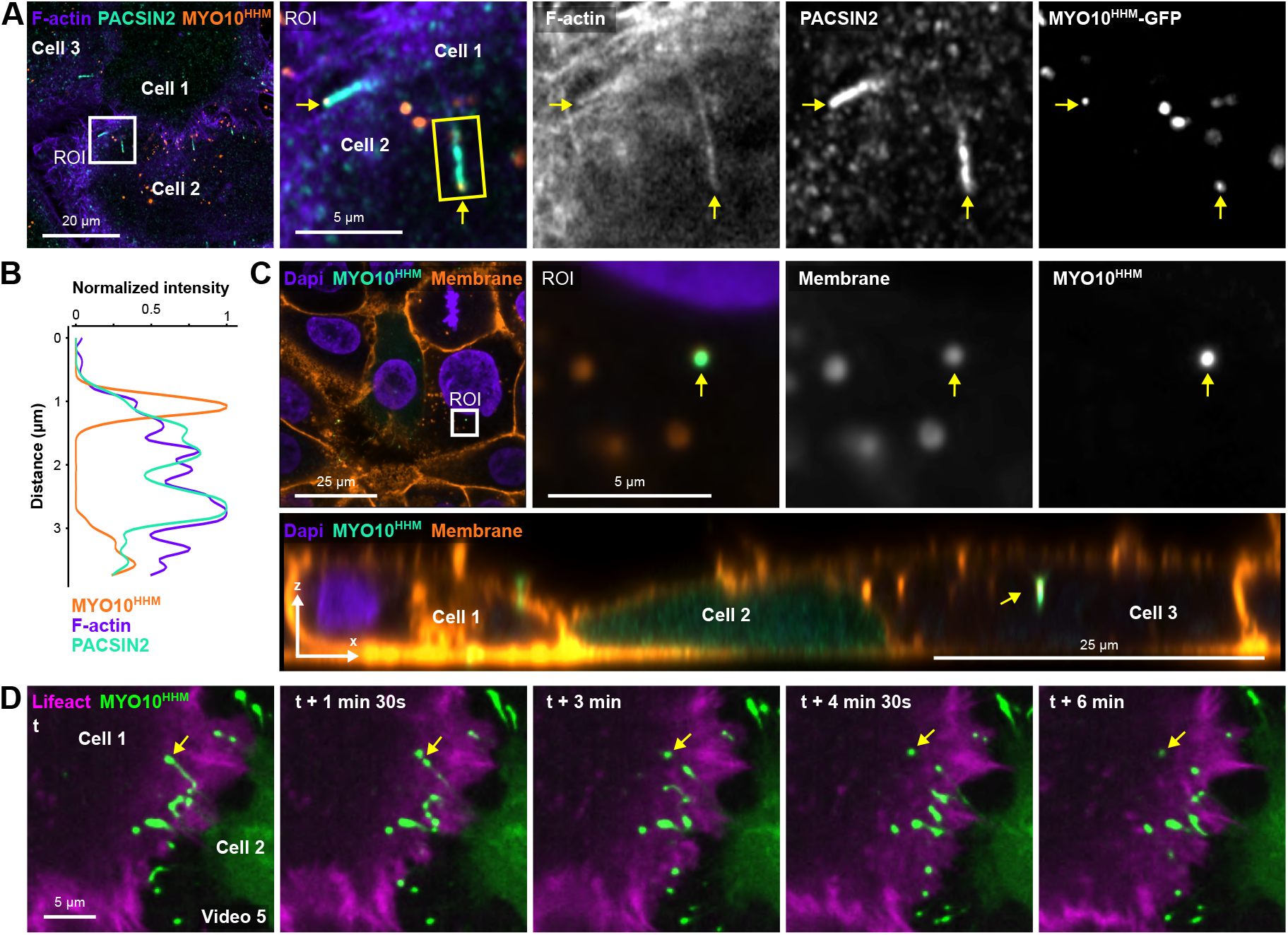
Intercellular filopodia tips can be internalised by neighbouring cells. (**A, B**) DCIS.com cells expressing MYO10^HMM^-GFP were cultured to form a monolayer, fixed, stained for endogenous PACSIN2, and imaged using Airyscan confocal microscopy. (**A**) A representative image is displayed. The yellow arrows highlight two sites of filopodia impact. (**B**) Example line intensity profiles of a filopodia impact site from the filopodia tip to the base. The yellow square highlights the filopodia used for the line-intensity profile (**C**) DCIS.com parental cells (cell 1 and cell 3) were co-cultured with DCIS.com MYO10^HMM^-GFP cells (cell 2), fixed, stained using fluorescently labelled Concanavalin A, and imaged using Airyscan confocal microscopy. Representative xy and xz planes are displayed. The yellow arrow highlights an internalised filopodia tip. (**D**) DCIS.com lifeact-RFP cells (cell 1) were co-cultured with DCIS.com MYO10^HMM^-GFP cells (cell 2) and imaged live using spinning disk confocal microscopy. The yellow arrows highlight the internalisation of filopodia tips in a lifeact-RFP-positive cell. See also **Video 5**. The raw images used to make this figure have been archived on Zenodo (Grobe et al., 2026).

To determine whether intercellular filopodia tips can be transferred between cells, we mixed DCIS.com MYO10^HMM^-GFP cells with DCIS.com lifeact-RFP cells. Remarkably, MYO10^HMM^-GFP-positive puncta were often found within lifeact-RFP-expressing cells, indicating that filopodia tips can be internalised by neighbouring cells (Fig. S7B). High-resolution 3D imaging confirmed the intracellular localisation of these puncta, which colocalise with cellular membranes (Fig. 4C). Finally, using live-cell imaging, we observed the active internalisation of MYO10^HMM^-GFP-labelled filopodia tips into neighbouring cells (Fig. 4D and Video 7).

Collectively, these findings demonstrate that tips of intercellular filopodia can be internalised by neighbouring cells, revealing a previously unrecognised mechanism of direct intercellular exchange and communication.

### Caveolin and dynamin are recruited to PACSIN2-positive filopodia impact sites

Having discovered that filopodia tips can be transendocytosed at filopodia impact sites, we next sought to identify the endocytic machinery recruited to these sites. We investigated whether canonical endocytic components, including markers of clathrin-mediated endocytosis (Clathrin Light Chain B, CLTB), caveolar endocytosis (caveolin-1, CAV1), and clathrin-independent carriers such as Arf-1 (ARF1) and insulin receptor substrate 53 (IRSp53, BAIAP2) (Kumari and Mayor, 2008; Sathe et al., 2018), accumulate at PACSIN2 fingers.

DCIS.com cells were transfected with GFP-tagged CLTB, CAV1, ARF1 or BAIAP2 and stained for PACSIN2 and F-actin. High-resolution Airyscan imaging showed a distinct and specific enrichment of CAV1 at filopodia impact sites (Fig. 5A). Conversely, CLTB, ARF1, and BAIAP2 exhibited little to no accumulation at these sites (Fig. 5B-D), suggesting that filopodia internalisation is unlikely to rely on clathrin-mediated endocytosis or the CLIC pathway. Live-cell imaging further confirmed the preferred association of CAV1 with filopodia impact sites (Video 8).

**Fig. 5.**
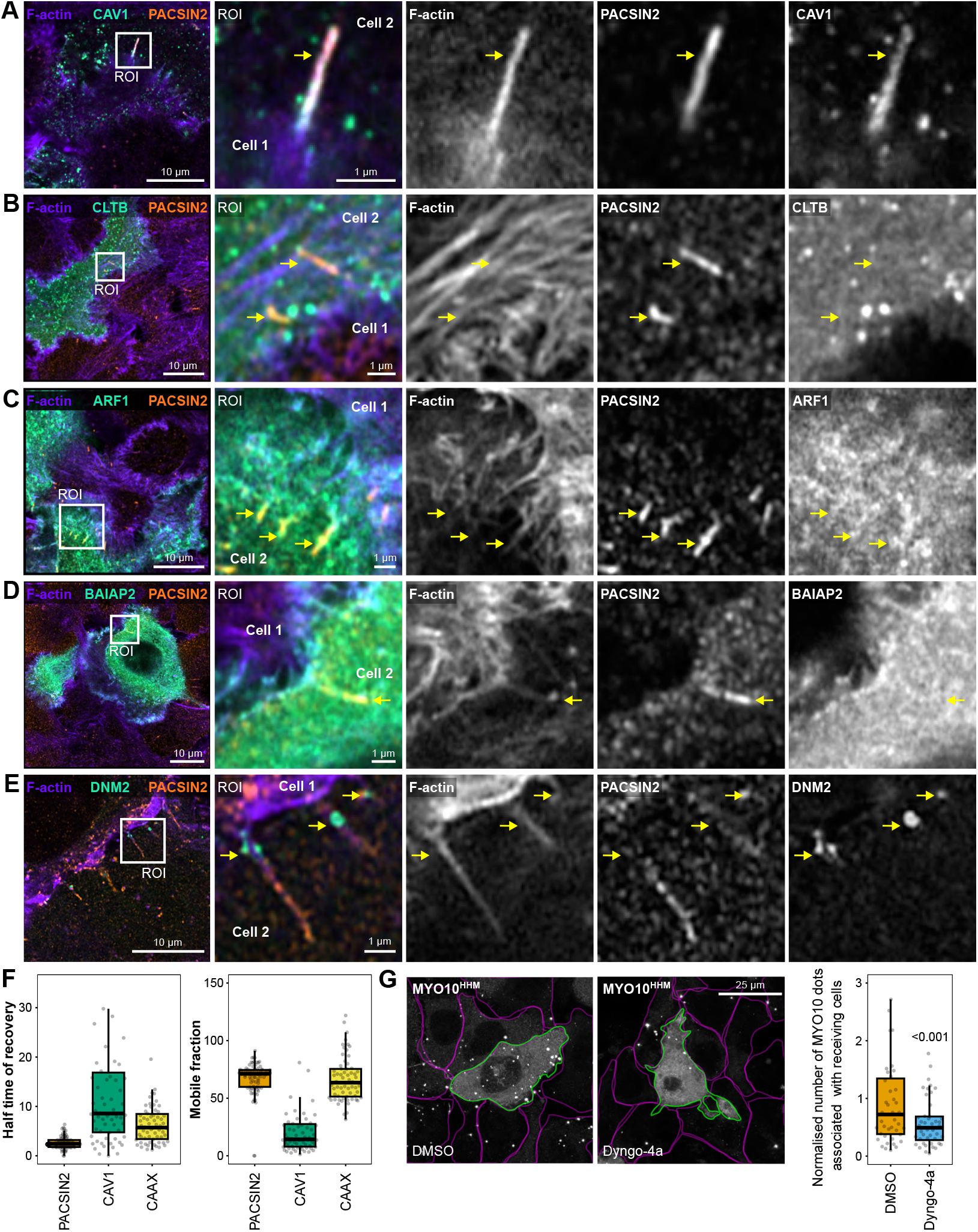
Caveolin and dynamin are selectively recruited to PACSIN2-positive filopodia impact sites. (**A-E**) DCIS.com parental cells were mixed with DCIS.com cells expressing GFP-tagged caveolin-1 (CAV1; **A**), clathrin light chain B (CLTB; **B**), Arf-1 (ARF1, **C**), IRSp53 (BAIAP2, **D**), or dynamin-2 (DNM2; **E**). Cells were grown as monolayers, fixed, stained for F-actin and endogenous PACSIN2, and imaged using Airyscan confocal microscopy. Representative images are shown. White squares indicate regions of interest that are magnified. Yellow arrows highlight filopodia impact sites. (**F**) FRAP analysis of GFP-tagged PACSIN2, CAV1, and CAAX at filopodia impact sites. Half-time of recovery and mobile fraction were quantified (3 biological replicates, > 66 filopodia impact sites). (**G**) DCIS.com parental cells (highlighted in magenta) were mixed with DCIS.com cells expressing MYO10^HMM^-GFP (highlighted in green). After 24 hours, cells were incubated with dynamin inhibitors Dyngo-4a or DMSO (10 µM, 24-hour treatment) before fixation. Cells were imaged using an Airyscan confocal microscope. Representative maximum intensity projections are displayed. MYO10^HMM^-GFP positive cells are outlined in green, while neighbouring cells are outlined in magenta. The number of MYO10^HMM^ dots associated with receiving cells was then quantified per field of view (see Methods, 3 biological replicates, >43 fields of view). Quantification was conducted per field of view, excluding the MYO10^HMM^-GFP-expressing cell; only MYO10^HMM^ dots associated with neighbouring parental cells were analysed (see Methods). Data are presented as boxplots, with whiskers extending from the 10th to the 90th percentiles. Boxes indicate the interquartile range, and the central line denotes the median. Data points outside the whiskers are shown as individual dots. The raw numerical values and images used to generate this figure have been archived on Zenodo (Grobe et al., 2026).

Since membrane scission is typically mediated by dynamin, we then examined the localisation of dynamin-2 (DNM2). DNM2-GFP was frequently recruited to filopodia impact sites in DCIS.com cells, where DNM2 accumulated either at the base of the filopodium or along the length of the impact site (Fig. 5E). This localisation pattern supports a role for DNM2 in the internalisation of filopodia tips.

To understand the dynamics of protein recruitment at filopodia impact sites, we conducted fluorescence recovery after photo-bleaching (FRAP) experiments. We compared the recovery kinetics of PACSIN2, CAV1, and a generic membrane marker (CAAX-GFP) at individual filopodia impact sites (Fig. 5F). PACSIN2 recovered more rapidly, with a shorter recovery half-time than CAAX-GFP, indicating highly dynamic exchange at these sites. In contrast, CAV1 exhibited a very slow recovery and a low mobile fraction, consistent with CAV1 being largely preassembled and stably associated with the membrane at these locations. Notably, PACSIN2 and CAAX-GFP had comparable mobile fractions, supporting a model in which PACSIN2 is dynamically recruited to pre-existing membrane domains rich in CAV1.

Finally, to test whether dynamin activity is required for filopodia tip exchange, we treated mixed monolayers of DCIS.com parental cells and DCIS.com MYO10^HMM^–GFP cells with the dynamin inhibitor Dyngo-4a and quantified MYO10^HMM^- positive puncta associated with recipient cells (Fig. 5G). Dyngo-4a significantly reduced the number of transferred MYO10^HMM^ puncta compared with DMSO-treated controls, indicating that dynamin activity promotes efficient filopodia tip internalisation.

Together, these data support the conclusion that filopodia transendocytosis takes place at specialised PACSIN2-positive impact sites that are enriched for CAV1 and DNM2.

### Internalised filopodia tips localise near ER tubules and can fuse with lysosomes

To investigate the fate of filopodia tips after internalisation, we performed correlative Airyscan fluorescence microscopy and focused ion beam-scanning electron microscopy (FIB-SEM) on mixed populations of DCIS.com cells expressing MYO10^HMM^-GFP and parental cells (Fig. 6A).

**Fig. 6.**
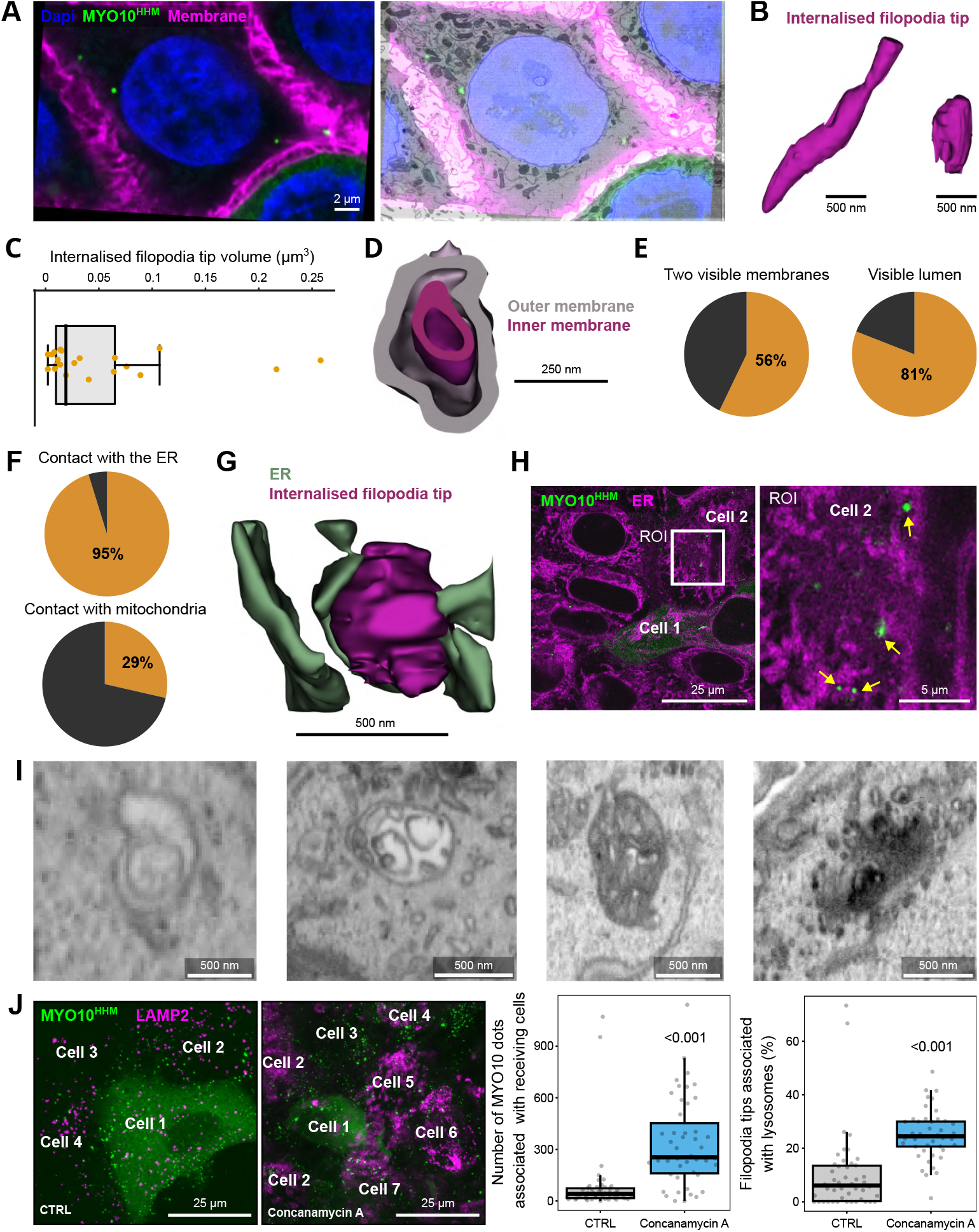
Internalised filopodia tips localise near ER tubules and can fuse with lysosomes. (**A-I**) DCIS.com cells expressing MYO10^HMM^-GFP were mixed with parental cells and grown as a monolayer. After 24 h, cells were fixed, labelled to visualise the plasma membrane and nuclei (DAPI), and imaged using high-resolution Airyscan fluorescence microscopy and focused-ion-beam scanning electron microscopy (FIB-SEM). (**A**) Representative registration and alignment of the fluorescence and FIB-SEM datasets. (**B**) Examples of internalised MYO10-positive filopodia tips identified and manually segmented from the FIB-SEM volume. (**C**) Quantification of the volume of internalised filopodia tips based on manual segmentation (n = 47 internalised filopodia tips, two biological replicates). (**D**) Representative FIB-SEM image showing an internalised MYO10-positive filopodia tip displaying a double-membrane structure. (**E**) Pie chart quantifying the proportion of internalised filopodia tips exhibiting two clearly visible membranes and a discernible lumen (n = 47 internalised filopodia tips, two biological replicates). (**F**) Quantification of the percentage of internalised filopodia tips in contact with endoplasmic reticulum (ER) tubules or mitochondria, as assessed from segmented FIB-SEM volumes (n = 47 internalised filopodia tips, two biological replicates). (**G**) Three-dimensional rendering of an internalised filopodia tip (magenta) in close association with ER tubules (green). (**H**) High-resolution Airyscan image showing an internalised filopodia tip in proximity to ER structures. (**I**) Representative electron microscopy images illustrating the ultrastructural diversity of internalised MYO10-positive filopodia tips. (**J**) DCIS.com cells expressing MYO10^HMM^-GFP were mixed with parental cells and cultured as a monolayer for 24 hours. Cells were then treated with concanamycin A (125 nM) for 24 hours. Subsequently, cells were fixed, stained for LAMP2, and imaged using an Airyscan confocal microscope. Representative images and quantification of the impact of lysosomal inhibition on the number of MYO10^HMM^ dots associated with recipient cells, as well as the percentage of MYO10^HMM^ dots overlapping with LAMP2 staining, are shown (4 biological replicates, 47 fields of view). Quantification was carried out per field of view, excluding the MYO10^HMM^-GFP-expressing cell; only MYO10^HMM^ dots associated with neighbouring parental cells were analysed (see Methods). (**C, J**) Data are presented as boxplots, with whiskers extending from the 10th to the 90th percentiles. Boxes indicate the interquartile range, and the central line denotes the median. Data points outside the whiskers are shown as individual dots. The raw images used to generate this figure have been archived on Zenodo (Grobe et al., 2026).

Registering fluorescence and EM datasets enabled us to identify internalised MYO10-positive filopodia tips within neighbouring cells, which were then analysed at ultrastructural resolution (Fig. S8A). Manual segmentation of EM structures corresponding to MYO10^HMM^-GFP signals allowed three-dimensional reconstruction and quantitative analysis of internalised filopodia tips (Fig. 6B). Volumetric measurements showed that most detached tips ranged from 0.01 to 0.1 µm^3^. However, some larger structures were also observed (Fig. 6B, C). In many cases, internalised filopodia tips were enclosed by two distinct membranes and contained a visible lumen, indicating the presence of opposed plasma membranes from donor and recipient cells (Fig. 6D, E).

To explore the intracellular environment of internalized filopodia tips, we examined their proximity to key organelles. Segmentation-based analysis revealed that 95% of internalised tips were directly in contact with endoplasmic reticulum tubules, while 29% touched mitochondria (Fig. 6F). 3D renderings further showed a close spatial relationship between the tips of internalised filopodia and ER membranes (Fig. 6G). These findings were confirmed by high-resolution Airyscan imaging, which consistently detected MYO10-positive puncta near ER-labelled structures in live cells (Fig. 6H).

Given the lysosome-like appearance of some internalised tips in the FIB-SEM data (Fig. 6I) and the known role of lysosomes in membrane and protein turnover, we next explored whether these tips could fuse with lysosomal compartments. High-resolution imaging of DCIS.com cells stained with SiR-lysosome showed partial colocalization between MYO10^HMM^- positive puncta and lysosomal markers, suggesting that some internalised tips are directed to lysosomes (Fig. S8B). To confirm this, lysosomal acidification was inhibited with concanamycin A. Inhibition for 24 hours significantly increased the number of internalised MYO10^HMM^-positive filopodia tips in cells neighbouring MYO10^HMM^-GFP positive cells and their accumulation within LAMP2-positive structures (Fig. 6J and Fig. S8C).

These data indicate that internalised filopodia tips frequently associate with ER tubules and can be transported to lysosomes, where they are likely degraded. However, it remains unclear whether components of internalised filopodia are partially recycled by recipient cells or solely targeted for lysosomal degradation.

### Filopodia-mediated trans-endocytosis happens during both homotypic and heterotypic cell-cell interactions

Filopodia-mediated trans-endocytosis is characterised by two main features: (i) formation of PACSIN2-positive “fingers” in the receiving cell at filopodia impact sites (Fig. 3), and (ii) internalisation of filopodia tips by neighbouring cells (Fig. 4). To determine whether this process occurs beyond DCIS.com monolayers, we examined additional cellular contexts.

We initially used U-2 OS cells (an osteosarcoma cell line), which produce few endogenous filopodia but robustly generate filopodia upon MYO10 overexpression (Jacquemet et al., 2019; Popović et al., 2025). When U-2 OS cells transiently expressing MYO10-mScarlet were mixed with those expressing PACSIN2-GFP, Airyscan imaging showed MYO10-positive filopodial puncta penetrating into neighbouring cells and initiating the formation of PACSIN2-positive fingers at the interface (Fig. 7A). To assess whether these structures could be internalised, we mixed U-2 OS cells expressing MYO10-mScarlet with those expressing the membrane marker LCK-GFP. In these co-cultures, MYO10-positive puncta were found inside MYO10-negative cells and colocalised with cellular membranes (Fig. 7B).

**Fig. 7.**
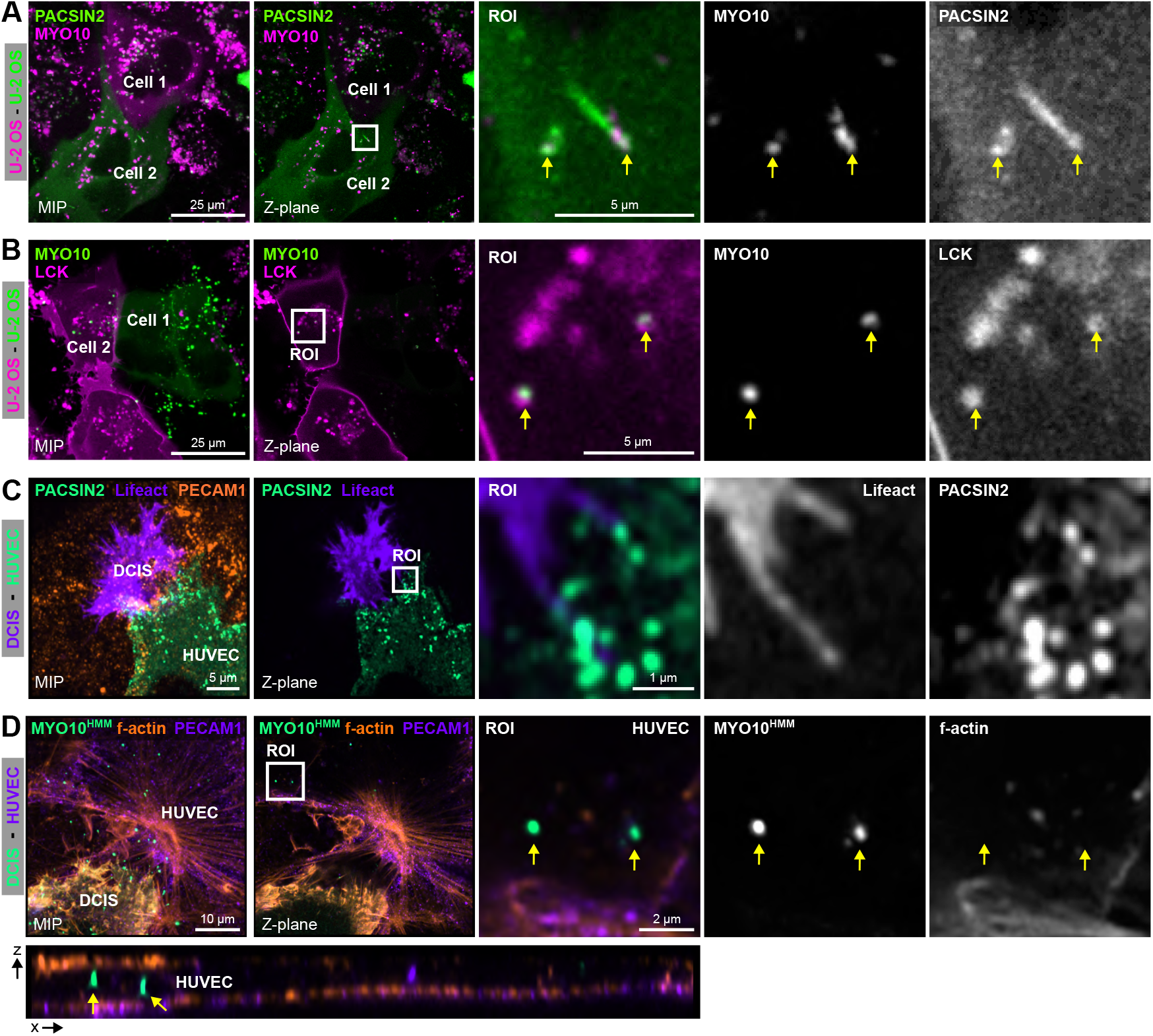
Filopodia-mediated trans-endocytosis happens during both homotypic and heterotypic cell-cell interactions. (**A**) U-2 OS cells transiently expressing MYO10-mScarlet were mixed with U-2 OS cells transiently expressing PACSIN2-GFP and imaged by Airyscan confocal microscopy. Representative images show PACSIN2-positive fingers forming at MYO10-positive filopodia impact sites. The white square indicates an ROI shown at higher magnification; yellow arrows mark representative impact sites. (**B**) U-2 OS cells transiently expressing MYO10-mScarlet were mixed with U-2 OS cells expressing LCK-GFP (membrane marker) and imaged by Airyscan confocal microscopy. The white square indicates an ROI shown at higher magnification; the yellow arrows highlight internalised MYO10-positive filopodia tips within MYO10-negative cells. (**C**) DCIS.com Lifeact-RFP cells were plated onto a HUVEC monolayer transiently expressing PACSIN2-GFP (mosaic), fixed, stained for PECAM1 (CD31), and imaged by Airyscan confocal microscopy. The white square indicates an ROI shown at higher magnification. (**D**) DCIS.com MYO10^HMM^-GFP cells were plated onto HUVEC monolayers, fixed, stained for PECAM1 (CD31) and F-actin, and imaged using Airyscan confocal microscopy. The white square marks an ROI shown at higher magnification. The yellow arrows indicate internalised MYO10-positive filopodia tips within MYO10-negative HUVEC cells. The same MYO10-positive filopodia tips are displayed in the xy and xz images. The raw images used to generate this figure have been archived on Zenodo (Grobe et al., 2026).

We then examined whether filopodia-mediated trans-endocytosis also occurs during heterotypic interactions relevant to cancer progression. We modelled cancer cell engagement with the endothelium by plating DCIS.com lifeact-RFP cells onto a monolayer of HUVECs transiently expressing PACSIN2-GFP in a mosaic pattern. As DCIS.com cells spread on and breached the endothelial monolayer, endothelial PACSIN2 assembled into finger-like structures around cancer cell filopodia at contact sites (Fig. 7C). Finally, when DCIS.com MYO10^HMM^-GFP cells were plated onto HUVEC monolayers, MYO10^HMM^-positive puncta were detected within endothelial cells (Fig. 7D), indicating that endothelial cells can internalise filopodia tips from invading cancer cells.

These results demonstrate that filopodia-mediated trans-endocytosis occurs not only in DCIS.com monolayers but also across different cellular systems, including both homotypic and heterotypic cell–cell interactions.

## Discussion

Here, we define filopodia-mediated trans-endocytosis as a contact-dependent process in which an intercellular filopodium penetrates a neighbouring cell to form a highly curved impact site on the recipient membrane, and some of these sites proceed to scission, thereby internalising part of the filopodium. The internalised filopodial material often associates with ER tubules and can be transported to lysosomes, indicating that this pathway links protrusion–cell contact with a specific intracellular trafficking route.

A notable feature of filopodia-mediated trans-endocytosis is the depth and shape of the impact sites. In epithelial monolayers, filopodial penetration can extend for micrometres into recipient cells, producing prominent invaginations. This raises an immediate mechanical question: How can a thin protrusion deform the recipient membrane–cortex complex so extensively? Actin polymerisation can generate force approximately 1 pN per filament (Footer et al., 2007), so a bundle of about 15 filaments per filopodium could theoretically produce around 15 pN, while a small group of tip-localised MYO10 motors could add several additional piconewtons (Fitz and Tyska, 2024; Homma and Ikebe, 2005), placing plausible filopodial pushing forces in the tens of pn range, though likely as upper estimates. Supporting this, direct force measurements in differentiating neurons suggest a single filopodium rarely exceeds about 3 pN of pushing force (Cojoc et al., 2007). In contrast, pulling a membrane tether, often used as a practical measure to overcome membrane bending and membrane–cortex attachment, typically requires forces of around 30 pN (Sun et al., 2005), which is close to or exceeds the upper limit of what a pushing filopodium might deliver. Collectively, these comparisons imply that the deep invagination found at filopodia impact sites probably does not indicate extraordinary protrusive power per se; rather, it likely depends on where the filopodium makes contact with the recipient surface and the local mechanical state of the cortex. Since cortical tension and membrane–cortex coupling are heterogeneous (Lieber et al., 2013), even slight local changes in actomyosin density, crosslinking, or membrane-cortex attachment could shift a region from resistance to permissiveness for filopodia-induced deformation. Consistent with this idea, CAV1 concentrates at impact sites, suggesting that caveolae, invaginated membrane reservoirs that can buffer tension (Sinha et al., 2011), may provide flexible entry points for filopodia. Supporting this, our FRAP measurements show rapid PACSIN2 exchange but slow CAV1 recovery, indicating that PACSIN2 is dynamically recruited to pre-existing caveolin-decorated filopodia impact sites. Overall, the depth and stability of filopodial impact sites likely reflect a balance between modest protrusive forces and locally adjustable recipient-cell mechanics. Future work combining cortex biophysics with high-resolution force measurements will be necessary to understand how filopodia overcome cortical resistance and how recipient cells sense and respond to this mechanical input.

At the molecular level, we observe strong recruitment of PACSIN2 and CAV1 to impact sites, forming dynamic “PACSIN2 fingers” in recipient cells. PACSIN2-positive tubular membranes have been described at endothelial adherens junctions, where PACSIN2 accumulates on the concave, high-curvature side of mechanically stressed, asymmetric junctions and helps stabilise VE-cadherin–based adhesion by limiting excessive cadherin internalisation. (Dorland et al., 2016; Malinova et al., 2021). However, the PACSIN2 fingers we describe differ in both location and behaviour as they originate below adherens junctions in epithelial monolayers and appear as transient, highly curved invaginations induced by filopodial impact, rather than junctional tension–driven remodelling. Notably, their architecture and protein composition resemble the membrane cups formed on recipient cells during Listeria cell-to-cell spread, where PACSIN2, caveolin, and dynamin-2 assemble at invasion sites to create a double-membrane compartment that internalises the protrusion (Sanderlin et al., 2019; Tijoriwalla et al., 2024). Overall, this suggests that PACSIN2 recruitment may act as a conserved curvature-handling module at cell–cell interfaces, which can either stabilise stressed junctions or facilitate controlled engulfment and material exchange, as exploited by pathogens such as Listeria by hijacking the trans-endocytic pathway documented here.

Our work also positions filopodia-mediated uptake within the broader landscape of trans-endocytosis and protrusion-driven exchange. Trans-endocytosis is traditionally described in receptor–ligand systems such as Notch–Delta and Eph–ephrin, where engagement across apposed membranes is coupled with internalisation to regulate signalling and remodel contacts (Marston et al., 2003; Qureshi et al., 2011; Valenzuela and Perez, 2020). Our findings reveal a specific trigger: a protrusion-driven, high-curvature impact point on the recipient membrane. This capture-and-scission process we describe here also differs from other protrusion-based exchanges. For instance, tunnelling nanotubes create longer-lasting, open channels for long-range cytoplasmic transfer (Ljubojevic et al., 2021). Cytonemes primarily serve as targeted signalling extensions that transport ligand–receptor complexes over long distances (Ma et al., 2025). We therefore propose that filopodia-mediated trans-endocytosis functions as a short-range, geometry-dependent exchange pathway, particularly suited to dense tissues, where tightly opposed, dynamic membranes may favour episodic uptake events over more stable intercellular tunnels.

A key unresolved issue is the nature of the cargo transferred during filopodia-mediated trans-endocytosis. Filopodia tips are not passive membrane extensions; they are highly organised subcellular compartments whose composition is shaped by diffusion, selective retention, and by motor-driven delivery of cargo (Jacquemet et al., 2019; Berg and Cheney, 2002; Hirano et al., 2011; Miihkinen et al., 2021; Popović et al., 2023). In this context, the filopodial tip can function as a specialised sorting microdomain (Bird et al., 2017), enriched with specific receptors, lipid species (Jacquemet et al., 2019; Mason et al., 2025), and cytoplasmic factors, including proteins and mRNA (Costa et al., 2020). Supporting the idea that filopodia can generate biologically active material for other cells, scission of filopodia by I-BAR protein in single cells has been shown to produce large extracellular vesicles that modulate the migration of recipient cells (Nishimura et al., 2021). Additionally, although filopodia are typically devoid of organelles, mitochondria have been reported to enter filopodia via MYO19 (Marlar-Pavey et al., 2025; Shneyer et al., 2016), indicating that the transferable repertoire may extend beyond membranes and cytoplasmic factors under certain conditions (Shneyer et al., 2017). Future work will focus on identifying the molecular composition of internalised filopodial structures, which will be vital to understanding the functional implications for recipient cells.

Finally, our trafficking observations suggest that degradation is common, with internalised filopodia tips building up in lysosomal compartments, especially when lysosomal function is inhibited, which supports delivery to degradative pathways. Furthermore, we did not observe direct reuse of transferred fluorescent proteins in newly formed filopodia. However, this does not rule out the possibility that some material follows alternative routes or exerts influences on cellular behaviour prior to degradation. Many receptors enriched in filopodia can signal from endosomal or late endosomal/lysosomal compartments (Valenzuela and Perez, 2020; Alanko et al., 2015; Vieira et al., 1996), raising the intriguing possibility that internalised filopodial material could support spatially confined signalling in recipient cells; how such signalling would occur within double membranes or multivesicular bodies remains unresolved.

Although this study offers a descriptive framework for filopodia-mediated trans-endocytosis, a major limitation is the absence of a perturbation that specifically inhibits the process without broadly affecting filopodia formation or cell viability. Consequently, we are currently unable to assess the influence of filopodia-mediated trans-endocytosis on collec-tive behaviour. Future priorities include identifying factors that determine whether an impact site undergoes scission or retraction, characterising the molecular payload and fate of internalised filopodial material, and investigating how filopodia-mediated trans-endocytosis influences tissue organisation in both physiological and pathological contexts.

## Methods

### Cells

MCF10 DCIS.com cell lines (DCIS.com; invasive T24 c-Ha-ras oncogene-transfected breast epithelial cells (Miller et al., 2000)) and immortalised normal breast epithelial cells (MCF10A) were cultured in a 1:1 mix of DMEM (Merck) and F12 (Merck) supplemented with 5% horse serum (GIBCO BRL, Cat Number: 16050122), 20 ng/ml human EGF (Merck, Cat Number: E9644), 0.5 mg/ml hydrocortisone (Merck, Cat Number: H0888-1G), 100 ng/ml cholera toxin (Merck, Cat Number: C8052-1MG), 10 mg/ml insulin (Merck, Cat Number: I9278-5ML), and 1% (vol/vol) penicillin/streptomycin (Merck, Cat Number: P0781-100ML) at 37°C, 5% CO_2_. MCF10A cells (CVCL 0598) used for spheroid generation were cultured in MEBM® Basal Medium (Lonza, CC-3151) supplemented with MEGM® SingleQuots® Supplements (Lonza, CC-4136).

T-47D luminal breast cancer cells (CVCL 0553) were cultured in Dulbecco’s Modified Eagle’s Medium (DMEM) supplemented with 10% fetal bovine serum (Gibco, A5256701) and 1% Penicillin–Streptomycin–Glutamine (Gibco, 10378016). All cell lines were maintained at 37°C in a humidified atmosphere containing 5% CO_2_. HEK293FT cells were cultured in DMEM HG, supplemented with 10% fetal bovine serum (FBS) (Biowest, S1860), 1% L-glutamine, and 1% penicillin-streptomycin. U-2 OS (human bone osteosarcoma epithelial cells) were cultured in DMEM (Dulbecco’s Modified Eagle’s Medium; Sigma, D1152) supplemented with 10% FBS (Biowest, S1860). All cell lines were routinely tested for mycoplasma infections and found to be free of mycoplasma.

DCIS.com cells were provided by J.F. Marshall (Barts Cancer Institute, Queen Mary University of London, London, England, UK). HEK293FT (CRL-1573), MCF10A (CRL-10317), and T-47D cells (HTB-133) were provided by the ATCC. U-2 OS cells were obtained from DSMZ (Deutsche Sammlung von Mikroorganismen und Zellkulturen, Braunschweig, Germany, ACC 785).

Human Umbilical Vein Endothelial Cells (HUVECs) (PromoCell, C-12203) were cultured in ready-to-use Endothelial Cell Growth Medium (ECGM) (PromoCell, C-22010 and C-39215), supplemented with 1% penicillin-streptomycin (Sigma-Aldrich, P0781). HUVECs from “P0” (commercial vial) were expanded to a P3 stock and stored at -80°C until used to standardise the experimental replicates. Vials were thawed, and the cells were transferred to a 10 cm culture dish for at least 2 days.

### Patient samples

Human breast tissue samples were collected from patients undergoing surgery for DCIS at the Department of Plastic and General Surgery at Turku University Hospital (Turku, Finland). All tissues were donated voluntarily with written informed consent (ethical approval ETKM 23/2018; amendments 20.3.2018, 19.2.2019, and 17.1.2023). Tissue samples were obtained by a pathologist after diagnostic preparation (see @tbl:patient_*s*_*amples*).

### Antibodies and reagents

The primary antibodies used in this study for immunofluorescence include anti-Pacsin-2 (1:100, PAB3301, Abnova Corporation), anti-LAMP-2 (1:100, DSHB, H4B4), and anti-CD31 (PECAM1) (Invitrogen, 370700). The secondary antibodies used at a 1:400 dilution for immunofluorescence, all sourced from Invitrogen, include Alexa Fluor 647-conjugated anti-mouse IgG (A21235), Alexa Fluor 488-conjugated anti-mouse IgG (A11001), Alexa Fluor 568-conjugated anti-mouse IgG (A10037), Alexa Fluor 647-conjugated anti-rabbit IgG (A21244), Alexa Fluor 488-conjugated anti-rabbit IgG (A11008), and Alexa Fluor 568-conjugated anti-rabbit IgG (A10042). Additional reagents used in this study include wheat germ agglutinin (WGA, ThermoFisher Scientific, W32466), ER-Tracker™ Red (ThermoFisher Scientific, E34250), SiR-lysosome (Spirochrome, SC012), Dyngo4a (Hydroxy-Dynasore) (Abcam, AB120689), DAPI (ThermoFisher Scientific, D1306), Alexa Fluor 647-conjugated phalloidin (Invitrogen, A30107), Poly-D-Lysine (Gibco™, A3890401), Alexa Fluor 647-conjugated Concanavalin A (ThermoFisher Scientific, C21421), Concanamycin A (Santa Cruz Biotechnology, sc-202111A), and fibronectin (Sigma-Aldrich, 341631). The fibrillar collagen I used for circular invasion assays was PureCol EZ Gel (Advanced BioMatrix, 5074). The Cultrex used for spheroid growth was obtained from Bio-Techne (3532-005-02), and the Methylcellulose for low-adhesion culture was obtained from Thermo Scientific (182312500).

### Plasmids

MYO10-mScarlet was previously generated and is available from Addgene (plasmid 145179) (Jacquemet et al., 2019). pCDH-lifeact-mRFP was a gift from P. Caswell (University of Manchester, UK). Arf1-EGFP was a gift from Marci Scidmore (Addgene plasmid 49578) (Moorhead et al., 2010). Lck-mScarlet-I was a gift from Dorus Gadella (Addgene plasmid 98821) (Chertkova et al.). The Caveolin1-GFP and Dynamin2-GFP plasmids were kindly provided by Johanna Ivaska (University of Turku, FI). mEmerald-Clathrin-15 (CLTB) was a gift from Michael Davidson (Addgene plasmid 54040) (Fiolka et al., 2012). The IRSP53/BAIAP2-GFP construct was described previously (Jacquemet et al., 2019). CAAX-GFP was a gift from Gregory Giannone (Bordeaux University, FR).

The PACSIN2-GFP plasmid (pcDNA6.2-PACSIN2-N-EmGFP) was generated by the cloning service of the Genome Biology Unit core facility (Research Programs Unit, HiLIFE Helsinki Institute of Life Science, Faculty of Medicine, University of Helsinki, Biocenter Finland), through transferring the PACSIN2 entry clone from the ORFeome collaboration library into mEmerald destination vectors using the standard LR reaction protocol.

The original MYO10^HMM^-GFP sequence (covering amino acids 2-948 of full-length bovine MYO10) was kindly provided by Timothy Andrew Sanders (University of Chicago, USA). The MYO10^HMM^-GFP sequence was PCR-amplified using Phusion Hot Start II (ThermoFisher Scientific, F549L) with primers containing KpnI and Eco32I sites. The amplicon and pENTR2b entry vector (ThermoFisher Scientific, A10463) were digested with FastDigest KpnI (ThermoFisher Scientific, FD0524) and Eco32I (ThermoFisher Scientific, FD0303) (vector digest included Shrimp Alkaline Phosphatase, 78390500UN), gel-purified (NucleoSpin Gel and PCR Clean-up; Macherey-Nagel), ligated with T4 DNA ligase (NEB M0202S), and transformed into DH5α cells. pENTR2b-MYO10^HMM^-GFP was miniprepped (NucleoSpin Plasmid EasyPure; Macherey-Nagel) and verified by sequencing. The verified pENTR2b-MYO10^HMM^-GFP entry clone was subsequently subcloned into pLenti6.3/V5-DEST using a standard LR reaction protocol by the Genome Biology Unit (Research Programmes Unit, HiLIFE Helsinki Institute of Life Science, Faculty of Medicine, University of Helsinki, Biocenter Finland) to produce the pLenti6.3/V5-DEST-MYO10-HMM-GFP lentiviral expression vector.

Generated plasmids are being deposited on Addgene (https://www.addgene.org/Guillaume_Jacquemet/).

### Transfection and generation of cell lines

HUVECs were electroporated using the Neon Transfection System (ThermoFisher Scientific, MPK5000) with the 10 µL kit (MPK1025; Thermo Fisher Scientific) following the manufacturer’s instructions (150,000 cells and 1 µg plasmid DNA per transfection) and as previously described (Ball et al., 2024). Transfected cells were then plated and supplemented with 50,000 untransfected HUVECs per well to facilitate monolayer formation.

DCIS.com and U-2 OS cells were transfected with the indicated plasmids using Lipofectamine 3000 and the P3000 Enhancer Reagent (Thermo Fisher Scientific, L3000001) according to the manufacturer’s instructions. DCIS.com lifeact-RFP cells were generated previously (Jacquemet et al., 2017) through lentiviral transduction using pCDH-lifeact mRFP, psPAX2, and pMD2.G constructs. The DCIS.com lifeact-RFP shCTRL and shMYO10 lines were generated previously (Peuhu et al., 2022) using lentiviral particles containing a non-target control shRNA (Merck, Cat Number: SHC016V-1EA) or shRNA targeting human MYO10 (shMYO10 3, TRCN0000123087; shMYO10 4, TRCN0000123088), respectively. Transduced cells were selected using normal media supplemented with 2 mg.ml-1 of puromycin. DCIS.com lifeact-RFP shCTRL and shMYO10 lines were generated from single-cell clones obtained from the DCIS.com lifeact-RFP shMYO10 3 and shMYO10 4 cell lines. Four single-cell clones with normal MYO10 levels were pooled to create the shCTRL line, and four single-cell clones with very low MYO10 levels were pooled to create the shMYO10 line.

DCIS.com MYO10^HMM^-GFP cells were generated by lentiviral transduction. HEK293FT cells were utilised to produce lentiviral particles. Cells were transfected with a third-generation lentiviral packaging system: the envelope plas-mid pMD2.G (a gift from Didier Trono, Addgene 12259), packaging plasmids pMDLg/pRRE (a gift from Didier Trono, Addgene 12251), pRSV-Rev (a gift from Didier Trono, Addgene 12253) (Dull et al., 1998), and pLenti6.3/V5-DEST-MYO10^HMM^-GFP as described above. The plasmids were co-transfected into HEK293FT cells at a 1:22:15:50 ratio, using 50 µL of Lipofectamine 3000 (Thermo Fisher Scientific, 100022052) in 8 mL of Opti-MEM (Gibco, 31985070). Twenty hours post-transfection, the medium was replaced with the standard growth medium. Viral particles were collected after 36 hours, filtered through 0.45 µm filters, and stored in 1 mL aliquots at -80°C. Parental DCIS.com cells were transduced with the lentiviral supernatant, mixed in a 1:1 ratio with fresh culture medium, and supplemented with polybrene (1:1000). After two weeks, stably transduced DCIS.com cells were sorted using fluorescence-activated cell sorting (FACS) on a BD FACSAria II cell sorter to isolate populations expressing MYO10^HMM^-GFP.

### Circular invasion assay

5 × 10^4^ DCIS.com or MCF10A cells were plated in a single well of a culture-insert 2-well (ibidi, 81176) pre-inserted within a well of a µ-Slide 8-well (ibidi). After 24 hours, the culture-insert 2-well was removed, and a fibrillar collagen (PureCol EZ Gel) gel was cast. The gels were allowed to polymerise for 30 minutes at 37°C before the normal medium was added. Cells were left to invade for 24 hours before fixation or live imaging.

### Spheroid generation

Single-cell suspensions of MCF10A and T-47D cells were prepared during trypsinisation by gentle trituration with a 1 mL syringe fitted with a 30-gauge needle (2–3 passes). For low-adhesion culture, cells were mixed with pre-filtered 2% methylcellulose (Thermo Scientific, 182312500) prepared in the respective cell culture medium to a final concentration of 1%, then seeded at 3,000 cells per well in 6-well low-adhesion plates. For 3D culture, cells were pelleted, resuspended in undiluted Cultrex basement membrane extract (Bio-Techne, 3532-005-02), and seeded approximately 500 cells in 45 µL BME per well in 8-well chamber slides.

After 10 days of culture, low-adhesion spheroids were collected by centrifugation, and BME domes were detached from chamber slides using a scalpel blade and transferred to Eppendorf tubes. The spheroid pellets were washed with 0.1 M cacodylate buffer and fixed in a fixative containing 2.5% glutaraldehyde, 4% paraformaldehyde, 0.1 M cacodylate buffer, and 2 mM CaCl_2_ for 4 hours at room temperature. Samples were then washed twice in cacodylate buffer, embedded in low-melting-point agarose, and allowed to solidify on ice. The embedded samples were overlaid with cacodylate buffer and processed for resin embedding and ultrathin sectioning for transmission electron microscopy.

**Table 1.** Patient characteristics and tumour details.

| ID | age | BMI | Diagnosis | Tumour size (mm) | DCIS grade |
| --- | --- | --- | --- | --- | --- |
| 301 | 63 | 27.5 | DCIS (with a small invasive focus) | 90 (DCIS) | 2-3 |
| 302 | 64 | 21.2 | DCIS | 51 | 2-3 |
| 303 | 64 | 20.5 | DCIS (no tumour detected in samples) | 0 | NA |

### Generation of tumour xenografts

Tumour xenografts were previously described (Peuhu et al., 2022), and this study used existing tissue material. Briefly, for xenografts, 13 x 10^5^ DCIS.com lifeact-RFP cells were resuspended in 100 ml of a mixture of 50% Matrigel (diluted in PBS) before being injected subcutaneously in the flank of 6-7-week-old virgin female NOD.scid mice (Envigo). Tumour growth was monitored with a calliper 1-2 times per week. Mice were sacrificed 10 or 25 days post-inoculation as indicated, and the tumours were dissected. Immunofluorescence stainings

Unless otherwise specified, cells were fixed in 4% paraformaldehyde (Thermo Fisher Scientific, 28908) in PBS for 10 minutes at room temperature. For LAMP2 staining, cells were fixed in 2% paraformaldehyde. Primary antibodies were diluted in PBS containing 5% horse serum; for LAMP2, the antibody solution also included 0.05% saponin (Thermo Fisher Scientific, 11453098). Primary antibody incubations were performed overnight at 4°C, followed by incubation with species-appropriate secondary antibodies (1:400) for 1 hour at room temperature. Alexa Fluor 647-phalloidin (1:800) and DAPI (1:3000 from a 5 mg/mL stock) were added during the secondary antibody step. Samples were washed at least twice with PBS between steps and mounted in ProLong Glass (Invitrogen, P36984).

### Interaction of cancer cells with endothelial cells

HUVECs were seeded in fibronectin-coated (10 µg/mL; 341631, Sigma-Aldrich) glass-bottom µ-slide eight-well chambers (80807; Ibidi) and cultured for 72 hours to form a monolayer (Follain et al., 2026). DCIS.com cells stably expressing lifeact-RFP or MYO10^HMM^-GFP were added onto HUVEC monolayers and fixed after 1 hour with prewarmed 4% paraformaldehyde (28908; Thermo Fisher Scientific) in PBS for 10 minutes at 37°C. Coverslips were mounted in Vectashield (H-1000-10; BioNordika) and imaged in 3D using an Airyscan confocal microscope.

### Light microscopy setup

The spinning-disk confocal microscope used was a Marianas spinning-disk imaging system equipped with a Yokogawa CSU-W1 scanning unit mounted on an inverted Zeiss Axio Observer Z1 microscope and operated via SlideBook 6 (Intelligent Imaging Innovations, Inc.). Images were captured using either an Orca Flash 4 sCMOS camera (chip size 2,048 × 2,048; Hamamatsu Photonics) or an Evolve 512 EMCCD camera (chip size 512 × 512; Photometrics). The objective employed was a 100x oil-immersion objective (NA 1.4, Plan-Apochromat, M27).

The structured illumination microscope (SIM) used was DeltaVision OMX v4 (GE Healthcare Life Sciences) equipped with a 60x Plan-Apochromat objective lens with a 1.42 NA (immersion oil RI of 1.516), operated in SIM illumination mode (five phases x three rotations). Emitted light was collected using a front-illuminated sCMOS camera (pixel size 6.5 µm, readout speed 95 MHz; PCO AG), controlled by SoftWorx.

The Airyscan confocal microscope used was an LSM880 (Zeiss) equipped with an Airyscan detector (Carl Zeiss) and either a 40x water (NA 1.2) or 63x oil (NA 1.4) objective. The microscope was operated with Zen Black (2.3), with Airyscan in standard super-resolution mode.

The confocal microscope used in this study was a Stellaris 8 FALCON (Leica) with FCS and Resonant Scanner, employing either a 63x PL APO CS2 63x/1.40 OIL or HC PL APO CS2 86x/1.20 water objective. The microscope was operated with Leica’s LAS X Software (Version 5.3.0).

### Expansion microscopy

DCIS.com cells were seeded on fibronectin-coated coverslips (10 µg/mL; 341631, Sigma-Aldrich) and cultured for 24 hours. Cells were fixed in 4% paraformaldehyde (28908; Thermo Fisher Scientific) in PBS for 10 minutes at 37°C, permeabilised with 0.2% Triton X-100 for 10 minutes at room temperature, and stained for PACSIN2 using anti-PACSIN2 (PAB3301, Abnova Corporation; 1:100) in PBS containing 2% BSA for 1.5 hours at room temperature, followed by Alexa Fluor 488 anti-rabbit secondary antibody (A11008, Invitrogen; 1:200) for 1 hour. Samples were anchored with AcX (A20770, Invitrogen; 100 µg/mL) overnight at room temperature, and F-actin was labelled with Actin ExM-561 (Crometra; 1:20) for 1 hour immediately before gelation.

A tenfold expansion was performed as previously described (Damstra et al., 2022). Gels were polymerised for 1 hour at 37°C in PBS containing 1.1 M sodium acrylate (7446-81-3, AK Scientific), 2 M acrylamide (A9099, Sigma), 90 µg/mL N,N-methylenebisacrylamide (M7279, Sigma), 1.5 mg/mL APS (9592.3, Roth), and 1.5 mg/mL TEMED (T9281, Sigma). The gels were detached and digested with Proteinase K (P8107S, New England Biolabs; 0.8 U/mL in PBS) for 3 hours at room temperature, followed by disruption and DAPI staining in 5% SDS, 200 mM NaCl, 50 mM Tris pH 7.5 containing DAPI (D1306, Invitrogen; 1:1000) for 3 hours at 80° C. The gels were rinsed in 0.4 M NaCl, washed twice in PBS (30 minutes each), and expanded in water.

Expanded gels were mounted on poly-L-lysine-coated Ibidi glass-bottom dishes (81158; Ibidi; 0.1% w/v poly-L-lysine, A-005-C, Merck Life Science) and imaged in 3D using an Airyscan confocal microscope in super-resolution mode with a 63×/1.4 NA oil objective.

### Filopodia formation assays on fibronectin

DCIS.com MYO10^HMM^-GFP cells and parental DCIS.com cells were seeded for 2 hours on glass-bottom, 8-well plates coated with 10µg/mL fibronectin. The cells were then fixed with 4% PFA, washed with PBS, and stained with phalloidin. To count filopodia, images were captured using a spinning-disk confocal microscope with a 100× objective, and filopodia per cell were manually counted based on the MYO10 or F-actin signal. Filopodia density at cell-cell interfaces.

Airyscan z-stacks of F-actin–stained MCF10A and DCIS.com monolayers were analysed using Fiji (Schindelin et al., 2012). Maximum-intensity projections were generated and used for all subsequent quantification. To enhance the detection of thin protrusions, images were processed with background subtraction (rolling-ball radius: 50 px), followed by a bandpass filter (large structures: 40 px; small structures: 3 px). Images were then converted to 8-bit and analysed with the Ridge Detection plugin (line width: 5; high/low contrast: 100/10; sigma: 0.7; lower/upper threshold: 0.17/1.87; minimum line length: 3 px) (Steger, 1998). Ridge outputs were saved as ROIs in the ROI Manager, and ROIs representing clear non-filopodial features were excluded prior to quantification. Cell–cell interfaces were defined by manually tracing basolateral junctions on a smoothed actin image (background subtraction, Gaussian blur σ = 3, contrast enhancement). Junctions were traced using a freehand line tool with a fixed line width (10 px) and added to the ROI Manager. The total junction length per field of view was recorded and exported. Filopodia and junction ROIs were converted into label images and analysed with the DiAna plugin to identify filopodia ROIs adjacent to or overlapping with traced junctions (nearest-neighbour setting k = 1) (Gilles et al., 2017). Filopodia density at cell–cell contacts was calculated as the number of junction-associated filopodia normalised to the total traced junction length for each field of view.

### Live structured illumination microscopy

DCIS.com lifeact-RFP cells were cultured on high-tolerance glass-bottom dishes (MatTek Corporation, coverslip 1.5) until they reached confluence. The cells were then imaged in standard media using 3D SIM (5 phases, 3 rotations, 1 µm volume, 80 ms exposure, 1% 568 nm laser power). Raw images were subsequently processed with SoftWorx. To eliminate reconstruction noise and artefacts, the images were further processed with a custom-trained CSBDeep CARE 3D model (Weigert et al., 2018).

The CARE 3D model used to restore live SIM images was trained on the ZeroCostDL4Mic platform and has been described previously (Hidalgo-Cenalmor et al., 2024; von Chamier et al., 2021). The training dataset included 24 paired images (1024x1024x33 pixels, with pixel sizes x, y: 40 nm, z: 125 nm) and is available on Zenodo (https://zenodo.org/records/3713337). To generate this dataset, DCIS.com cells expressing lifeact-RFP were plated on high-tolerance glass-bottom dishes (MatTek Corporation, coverslip 1.5) and allowed to reach confluence. The cells were then fixed and permeabilised simultaneously using a solution of 4% (w/v) paraformaldehyde and 0.25% (v/v) Triton X-100 for 10 minutes. Afterwards, cells were washed with PBS, quenched with 1 M glycine for 30 minutes, and incubated with phalloidin-488 (1/200 in PBS; A12379; Thermo Fisher Scientific) at 4 °C overnight, then imaged. Prior to SIM imaging, the samples were washed 3 times in PBS and mounted in Vectashield (Vector Labs). In the training dataset, high-signal-to-noise-ratio images were obtained via phalloidin-488 staining, with acquisition parameters optimised for high-quality SIM images (50 ms exposure time, 10% laser power). Conversely, low-signal-to-noise-ratio images were acquired from the lifeact-RFP channel using settings more appropriate for live-cell imaging (100 ms exposure time, 1% laser power).

### Transmission electron microscopy

Patient samples and mouse tumour xenografts were fixed with 5% glutaraldehyde in s-collidine buffer at room temperature for 24 hours, post-fixed with 1% osmium tetroxide containing 1.5% potassium ferrocyanide, dehydrated through a series of increasing concentrations of ethanol and acetone, and gradually infiltrated into Fluka Epoxy Embedding Medium kit (45359). Thin sections were cut with an ultramicrotome to a thickness of 70 nm. The sections were stained with uranyl acetate and lead citrate. The sections were examined using a JEOL JEM-1400 Plus transmission electron microscope operated at 80 kV, equipped with an OSIS Quemesa 11 Mpix bottom-mounted digital camera. The spheroids in agarose blocks were then postfixated with 1% non-reduced osmium tetroxide in 0.1 M sodium cacodylate buffer, pH 7.4 (NaCac), for 1 hour and dehydrated through a series of increasing ethanol and acetone concentrations prior to gradual infiltration into epoxy (TAAB 812, Aldermaston, UK). To localise the spheroids, semithin 500-nm sections were cut and stained with Toluidine blue for light microscopy to guide trimming of TEM pyramids for spheroid location. Ultrathin 60-nm sections were cut and post-stained with uranyl acetate and lead citrate. The sections were examined using a JEOL JEM-1400 transmission electron microscope operated at 80 kV (Jeol Ltd., Tokyo, Japan), equipped with a bottom-mounted CCD camera (Orius SC 1000B, Gatan Inc., Pleasanton, CA). Cell-cell interface curvature was quantified using the Fiji plugin “Kappa” based on manual tracing of EM images

### Correlative focused ion beam electron microscopy

Cells were seeded into ibidi 2-well culture inserts (ibidi, 81176) placed at the centre of MatTek 35 mm dishes with gridded cover slips (P35G-1.5-14-CGRD MatTek Ashland, MA) coated with poly-L-lysine and fibronectin. Cells were fixed with 2% formaldehyde and 0.05% glutaraldehyde in 0.1 M HEPES supplemented with 2 mM CaCl_2_ and 2 mM MgCl_2_ for 25 minutes at room temperature, then stained with WGA (1:1000) and DAPI. Between each step, cells were washed three times with 0.1 M HEPES supplemented with 2 mM CaCl_2_ and 2 mM MgCl_2_.

Regions of interest were initially imaged using an Airyscan confocal microscope; the etched coordinates were documented with phase-contrast microscopy. After fluorescence imaging, cells were post-fixed with 2% glutaraldehyde in 0.1 M HEPES supplemented with 2 mM CaCl_2_ and 2 mM MgCl_2_ for 25 minutes at room temperature. The samples were washed and treated with 1% reduced osmium tetroxide in 0.1 M NaCac buffer supplemented with 2 mM CaCl_2_ on ice for 1 hour. This was followed by incubation in 1% aqueous thiocarbohydrazide (Sigma-Aldrich) for 10 minutes at room temperature, 1% unbuffered osmium tetroxide for 1 hour at room temperature, and 1% aqueous uranyl acetate at 4°C overnight. Between treatments, samples were rinsed five times in ion-exchanged water (each for 1 minute at room temperature). Before plastic embedding, samples were dehydrated through a graded ethanol series and then progressively infiltrated with Durcupan (ACM resin; Electron Microscopy Sciences) according to the manufacturer’s instructions. After polymerisation at 60°C for 48 hours, the embedded sample was detached from the coverslip by immersion in liquid nitrogen. The target area, identified by coordinates obtained from light microscopy, was excised, mounted on an aluminium pin with conductive glue, and coated with a thin layer of platinum (Quorum Q150TS, Quorum Technologies, Laughton, UK).

The correlated cells were imaged using a Zeiss FIB-SEM (Crossbeam 550 with Gemini 2 optics, Carl Zeiss Microscopy GmbH, Jena, Germany) at 5 nm pixel size with a 10 nm milling depth. The SEM micrographs were acquired with a 0.31 nA electron beam, 1.6 keV voltage, 1 µs dwell time, and 4× line averaging. Both backscattered and secondary electron signals were collected using inlens detectors. For milling new surfaces, the FIB operated at a beam current of 3 nA and an acceleration voltage of 30 kV. The dataset was collected and aligned using Zeiss Atlas 5 software with the 3D tomography module. FIB-SEM and Airyscan volumes were aligned using the eC-CLEM plugin (Paul-Gilloteaux et al., 2017) in icy (de Chaumont et al., 2012) using the default settings.

Segmentation of the FIB-SEM datasets was done manually using webKnossos (Boergens et al., 2017). For quantitative analyses, filopodia were manually traced in the FIB-SEM dataset using the ‘3D lines’ tool in Microscopy Image Browser (Belevich et al., 2016).

Segmented FIB-SEM volumes were rendered using 3D Slicer (Fedorov et al., 2012). Animations were created using the Animator module from the SlicerMorph extension (Rolfe et al., 2021) in 3D Slicer.

### PACSIN2 finger dynamics

DCIS.com Lifeact-RFP cells were transiently transfected with PACSIN2-GFP and plated to form a confluent monolayer. Live imaging was performed on a Zeiss LSM880 equipped with an Airyscan detector (standard super-resolution mode). Time-lapse sequences were captured as single optical sections (single z-plane) at 3-second intervals for 100 frames. PACSIN2-positive “fingers” were segmented in Fiji (Schindelin et al., 2012). Image stacks were first denoised using a median filter (radius 2 pixels) applied to the time series, then thresholded with the Shanbhag method (dark background) and converted to binary masks. Individual PACSIN2-positive objects were identified via Analyse Particles (area 0.05–22.28 µm^2^) and exported as labelled images for tracking. Object trajectories were reconstructed in Fiji with TrackMate v7 (Ershov et al., 2022). Tracking was performed on the labelled images using the Label Image Detector and linked with the LAP tracker (maximum linking distance 1 µm; maximum gap-closing distance 1 µm; maximum gap 1 frame). Track splitting was enabled with a splitting distance threshold of 0.5 µm. Track-level metrics were exported from TrackMate for further analysis and plotting.

### Fluorescence recovery after photobleaching (FRAP)

DCIS.com wild-type cells were transiently transfected with the indicated constructs, mixed with DCIS.com Lifeact-RFP cells, and plated to form a confluent monolayer. FRAP imaging was performed on a 3i Marianas spinning disk microscope. Impact sites were visually identified using the Lifeact-RFP signal. A single ROI was drawn over the impact site using the Slidebook ROI tool and designated for photobleaching. Images were analysed using the “FRAPanalysis.py” script in FIJI (downloaded from https://imagej.net/tutorials/analyzefrap-movies).

### Quantification of exchanged MYO10 puncta

Fluorescence images were processed in Fiji. Briefly, maximum-intensity projections were created. Images were cropped by 12 pixels at the borders to eliminate edge artefacts. MYO10^HMM^-positive puncta, representing filopodia tips, were analysed in relation to LAMP2-positive lysosomes. To exclude intracellular signals and filopodia tips still attached to the cell body, a single-cell ROI outlining the transfected cell was manually drawn on the MYO10^HMM^ (488 nm) projection and isotropically expanded by 5 µm. Pixel intensities within this expanded ROI were cleared, removing the cell body and proximal filopodia, leaving only regions spatially separated from the cell.

Puncta detection was conducted on the remaining extracellular area after median filtering (radius 1 px), using channel-specific detection parameters (MYO10^HMM^, 488 nm: intensity threshold 5, size 5, shape 14; LAMP2, 647 nm: intensity threshold 10, size 10, shape 35). Only MYO10^HMM^-positive filopodia tips outside the expanded cell boundary were retained. Label images were produced for each channel.

Colocalisation and nearest-neighbour proximity between MYO10^HMM^-positive filopodia tips and LAMP2-positive lysosomes were quantified using the DiAna plugin (Gilles et al., 2017).

### Quantification and statistical analysis

The code for conducting randomisation tests and t-tests is available on GitHub (https://github.com/CellMigrationLab/Plot-Stats). Randomisation tests used Cohen’s d as the effect-size metric, with 10,000 iterations per test. For t-tests, the data were assumed to be normally distributed, although this was not formally tested.

## Data availability

The raw microscopy images and the numerical data used to make the figures have been archived on Zenodo (Grobe et al., 2026).

The authors declare that the data supporting the findings of this study are available within the article and from the authors upon request. Any additional information needed to reanalyse the data reported in this paper is available from the corresponding authors.

## Manuscript preparation

Figures were prepared using Fiji and Inkscape. PlotsOfData (Postma and Goedhart, 2019). Volcano plots were generated with VolcaNoseR (Goedhart and Luijsterburg, 2020). SuperPlots were created using SuperPlotsOfData (Goedhart, 2021). GPT-5 (OpenAI) and Grammarly (Grammarly, Inc.) served as writing aids during manuscript preparation. The author also edited and validated all sections of the text. GPT-5 did not provide references. The PDF version of this manuscript was formatted with Rxiv-Maker (Saraiva et al., 2025).

## Author contributions. Conceptualisation

E.P., E.J., GJ. **Methodology**: H.G., M.R., S.G., MC.L., A.N., M.V., H.V., A.P., J.T., M.O., P.B., P.H., J.E., E.P., E.J., GJ. **Reagents**: P.B., P.H., E.J., G.J. **Formal Analysis**: H.G., M.R., S.G., MC.L., A.N., M.V., H.V., M.O., GJ. **Investigation**: H.G., M.R., S.G., MC. L., A.N., M.V., H.V., A.P., J.T., M.O., E.P., GJ. **Writing - Original Draft**: G.J. **Writing - Review and Editing**: Everyone. **Visualisation**: H.G., M.R., S.G., M.V., GJ. **Supervision**: G.J. **Funding Acquisition**: G.J.

### Acknowledgments

This study was funded by the Research Council of Finland (338537, 371287, and 374180 to G.J.), the Sigrid Juselius Foundation (to G.J.), the Cancer Society of Finland (Syöpäjärjestöt; to G.J.), and the Solutions for Health strategic funding for Åbo Akademi University (to G.J.). Additionally, this research was supported by the InFLAMES Flagships Programme of the Research Council of Finland (decision numbers: 337530, 337531, 357910, and 35791). G.J. is supported by the Finnish Cancer Institute (K. Albin Johansson Professorship).

Imaging was carried out at the Advanced Imaging Core Facility at Turku Bioscience Centre, the Electron Microscopy Core Facility at the Institute of Biomedicine, University of Turku, and the Electron Microscopy Unit at the Institute of Biotechnology, Helsinki Institute of Life Science. These facilities received support from Turku BioImaging, Helsinki Institute of Life Science, Biocenter Finland, and the Finnish Advanced Microscopy Node of Euro-BioImaging Finland (funded by the Research Council of Finland, FIRI 2023 grant decision numbers 359073 and 358879, and FIRI 2024 grant decision numbers 367582 and 367577).

## ABOUT THIS MANUSCRIPT

This work is licensed under CC BY 4.0.

## Supplementary Information

***Video 1. Live imaging of actin dynamics at the cell–cell interface*. Part A**: Live structured illumination microscopy (SIM) of confluent DCIS.com cells expressing lifeact-RFP displaying highly dynamic intercellular filopodia at basolateral cell–cell interfaces. A 1-µm-thick volume is shown as a maximum-intensity projection. Raw SIM data were reconstructed and denoised using CSBDeep CARE (see Methods). **Part B**: Live Airyscan imaging of confluent DCIS.com cells expressing Lifeact-RFP, illustrating intercellular filopodia dynamics at basolateral cell–cell interfaces. A single z-plane is displayed. In both parts, filopodia repeatedly protrude into intercellular gaps, bend, and retract during collective migration, continuously remodelling the contact zone.

***Video 2. FIB–SEM volume rendering of the DCIS*.*com cell–cell interface***. Focused ion beam–scanning electron microscopy (FIB–SEM) volume rendering of a DCIS.com epithelial monolayer showing extensive interdigitated cell–cell interfaces and frequent filopodia penetration events. Manual segmentation was performed in webKnossos, and rendering was carried out in 3D Slicer using SlicerMorph.

***Video 3. Filopodia penetration and impact-site architecture in FIB–SEM***. Orthogonal browsing through a representative FIB–SEM subvolume showing intercellular filopodia penetrating a neighbouring cell. Manual segmentation was performed in webKnossos, and rendering was carried out in 3D Slicer using SlicerMorph.

***Video 4. Additional impact-site configurations uncovered by FIB–SEM***. FIB–SEM subvolumes highlight distinctive impact-site structures, including (i) “filopodia caves” formed by multiple penetrations into a shared invagination, (ii) one impact site created by filopodia from two neighbouring cells, (iii) two impact sites created by a single filopodium through bifurcation, (iv) a cis impact site where a filopodium folds back and invaginates into its originating cell, and (v) curved and undulated filopodia. Manual segmentation was performed in webKnossos, and rendering was carried out in 3D Slicer using SlicerMorph.

***Video 5. Dynamics of PACSIN2 “fingers” at filopodia impact sites*. Part A**: Live Airyscan imaging of lifeact-RFP DCIS.com cells transiently expressing PACSIN2-GFP. Note the recruitment of PACSIN2 to filopodia impact sites. **Part B**: Live Airyscan imaging of a mix of lifeact-RFP DCIS.com with parental DCIS.com cells transiently expressing PACSIN2-GFP. Part C: Live Airyscan imaging of lifeact-RFP DCIS.com cells transiently expressing PACSIN2-GFP. Note the cleavage of the PACSIN2 finger. For all videos, a single Z-plane is displayed.

***Video 6. Expansion microscopy reveals PACSIN2 organisation at filopodia impact sites***. Three-dimensional Airyscan imaging of 10× expanded DCIS.com monolayers stained for endogenous PACSIN2 and F-actin, showing PACSIN2 forming a sheath-like structure around the penetrating actin-rich filopodium. Rendering was performed using Arivis Vision4D.

***Video 7. Filopodia tip transfer by trans-endocytosis***. Live imaging of co-cultured DCIS.com cells shows MYO10HMM-GFP tagging filopodia tips in donor cells and lifeact-RFP marking actin in neighbouring cells. The movie depicts a MYO10^HMM^- positive filopodial tip forming an impact site, followed by its scission and internalisation into the recipient cell, consistent with filopodia-mediated trans-endocytosis.

***Video 8. CAV1 at filopodia impact sites***. Live Airyscan imaging of lifeact-RFP DCIS.com cells transiently expressing CAV1-GFP (Caveolin-1). Note the presence of CAV1 at filopodia impact sites and its internalisation.

---

**Supplementary Figure S1.**
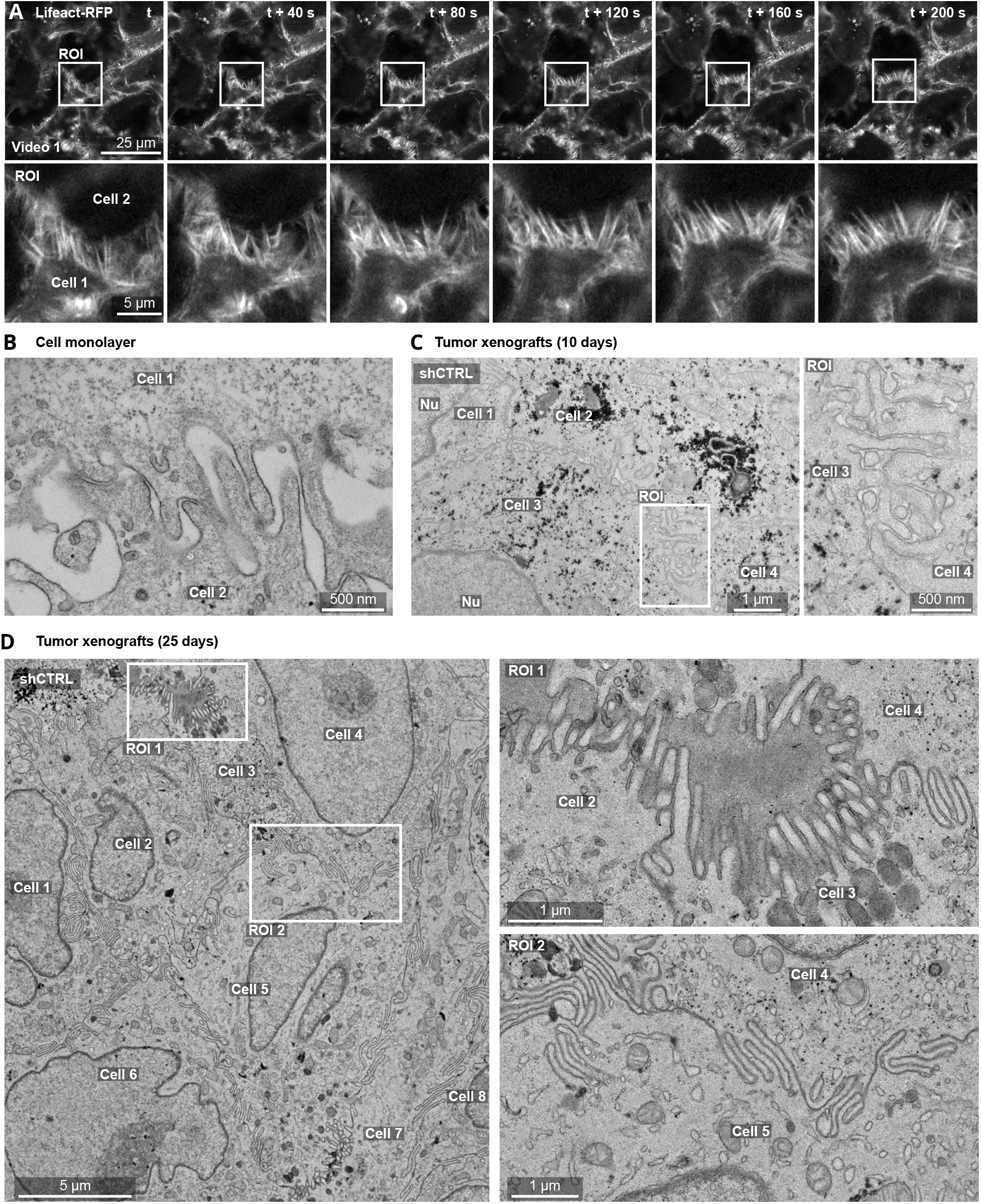
Intercellular filopodia are present in vitro and DCIS-like xenografts. (**A**) DCIS.com lifeact-RFP cells were seeded in circular invasion assays and left to invade into fibrillar collagen I for 24 hours. Cells were imaged live using Airyscan confocal microscopy. A representative single Z-plane showing filopodia on the lateral cell surfaces, with magnified ROIs, is shown (see also Video 1). Scale bars: main image, 25 µm; ROIs, 5 µm. (**B**) DCIS.com cell monolayers were fixed and imaged using electron microscopy (EM). A representative EM image illustrating intercellular filopodia is shown. Scale bar: main image, 500 nm. (**C, D**) 10-day-old (**C**) and 25-day-old (**D**) shCTRL DCIS-like xenografts were imaged using EM to visualise the cell-cell interface. Representative images and magnified ROIs are displayed. (**C**) Scale bars: main image, 1 µm; ROI, 500 nm. (**D**) Scale bars: main image, 5 µm; ROIs, 1 µm. The raw images used to make this figure have been archived on Zenodo (Grobe et al., 2026).

**Supplementary Figure S2.**
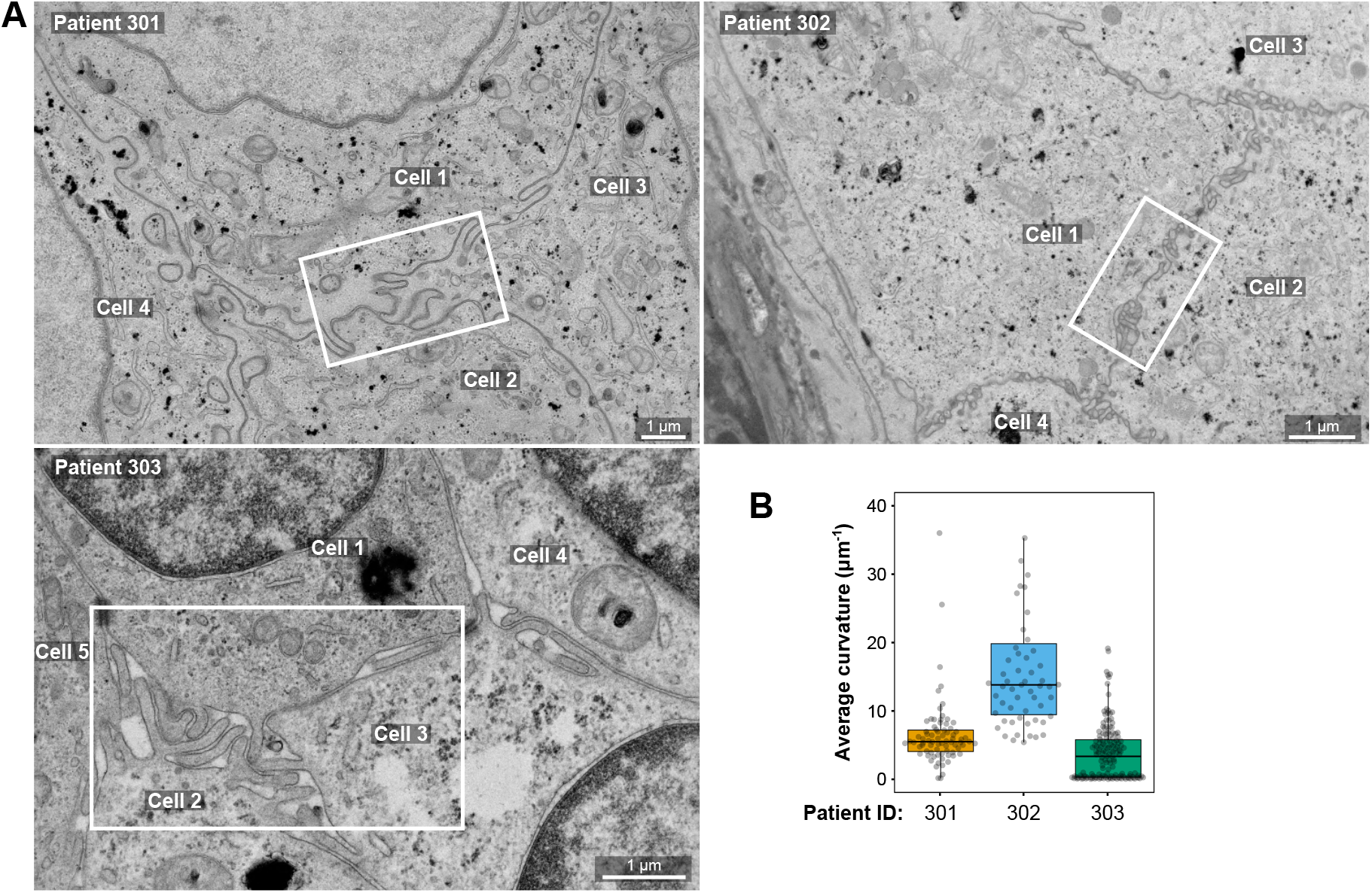
Intercellular filopodia are present in patient-derived DCIS lesions. (**A-B**) Patient-derived tissues processed and imaged by EM. Representative images showing examples of intercellular filopodia at cell-cell interfaces in each patient. White squares indicate the ROI displayed in Fig. 1G. Scale bars: 1 µm. (**B**) Quantification of cell-cell interface curvature using the Fiji plugin “Kappa” based on manual tracing of EM images (n > 59 FOV, 1 biological repeat). Results are shown as boxplots, with whiskers extending from the 10th to the 90th percentiles. The boxes indicate the interquartile range, and a line within each box marks the median. Data points outside the whiskers are shown as individual dots. The numerical data and the raw images used to make this figure have been archived on Zenodo (Grobe et al., 2026).

**Supplementary Figure S3.**
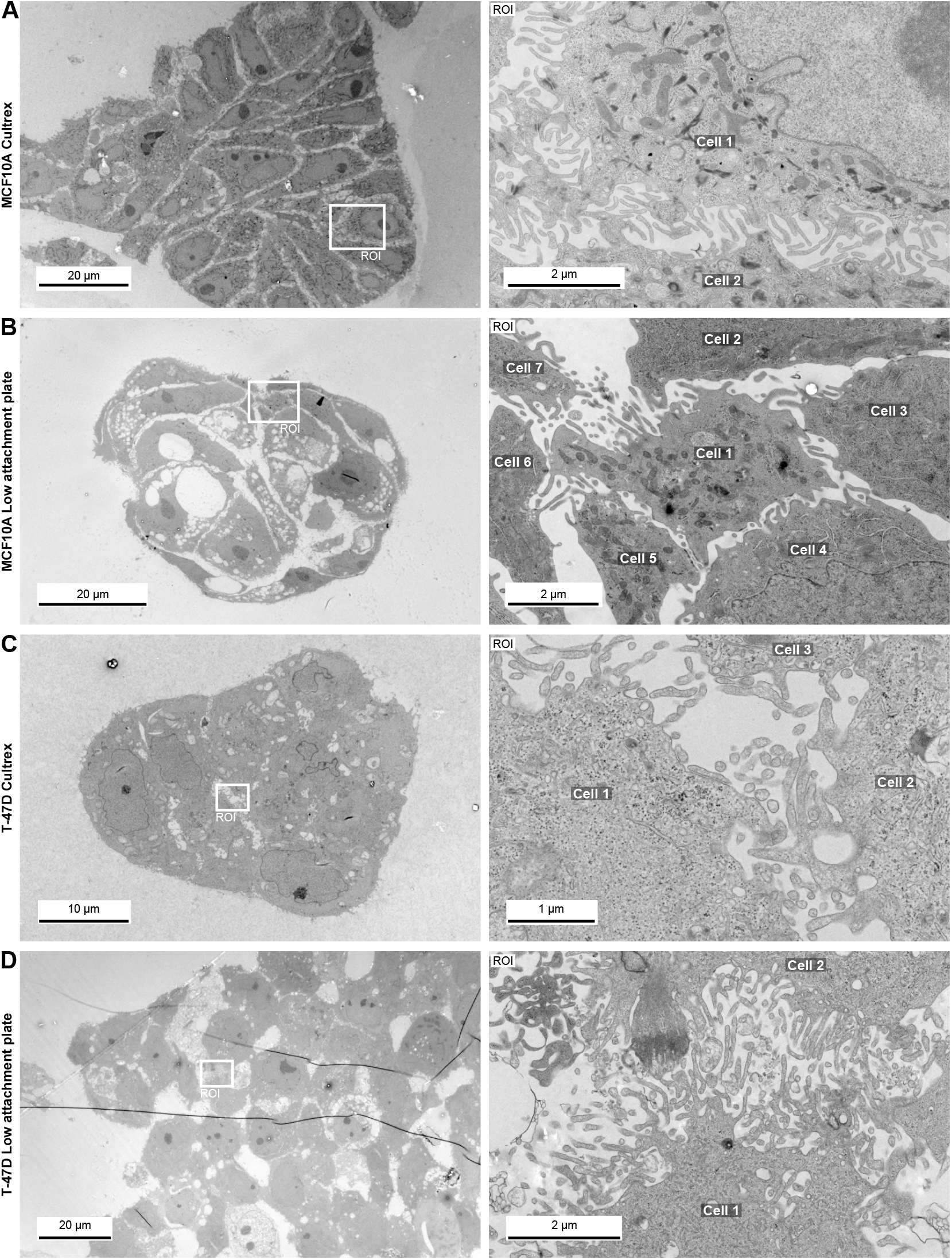
Intercellular filopodia are present in spheroids. (**A-D**) MCF10A (**A-B**) and T-47D (**C-D**) cells were grown as spheroids, either by embedding single cells in Cultrex (**A, C**) or by using low-attachment plates (**B, D**). Spheroids were then processed for transmission electron microscopy. For each condition, a representative low-magnification image of an entire spheroid and a corresponding high-magnification image focusing on cell-cell connections are shown. The raw images used to generate this figure have been archived on Zenodo (Grobe et al., 2026).

**Supplementary Figure S4.**
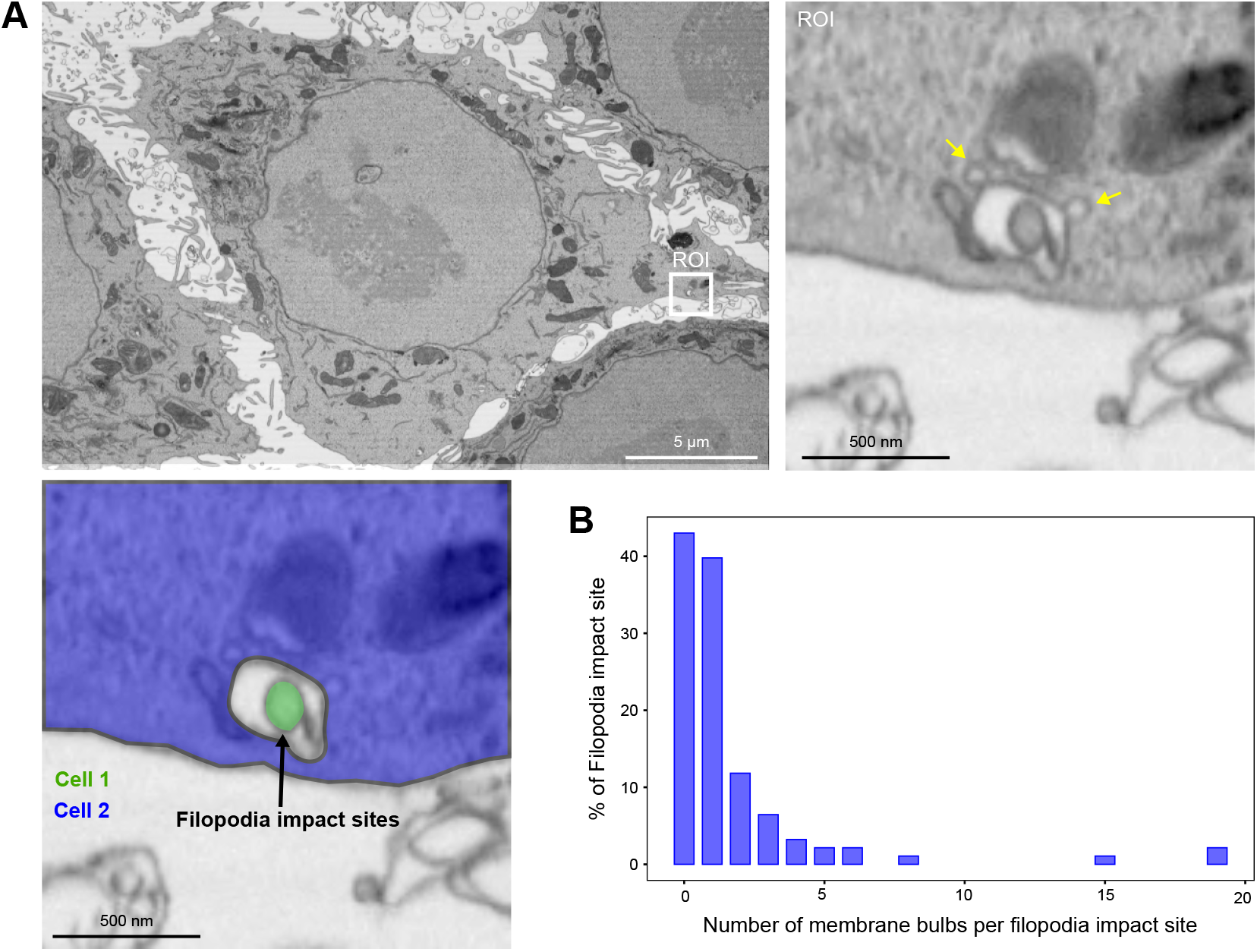
Membrane bulbs are often observed at filopodia impact sites. (**A**) Representative FIB-SEM image showing an ROI centred on a filopodia impact site, in which two membrane bulbs are highlighted (yellow arrows). The cellular origin of the filopodium is indicated and was manually assigned by tracing the protrusion throughout the FIB-SEM volume. Scale bars: main image, 5 µm; ROI, 500 nm. (**B**) Frequency histogram showing the number of membrane bulbs per filopodia impact site, quantified from two FIB-SEM volumes acquired in two independent experiments (n = 105 filopodia).

**Supplementary Figure S5.**
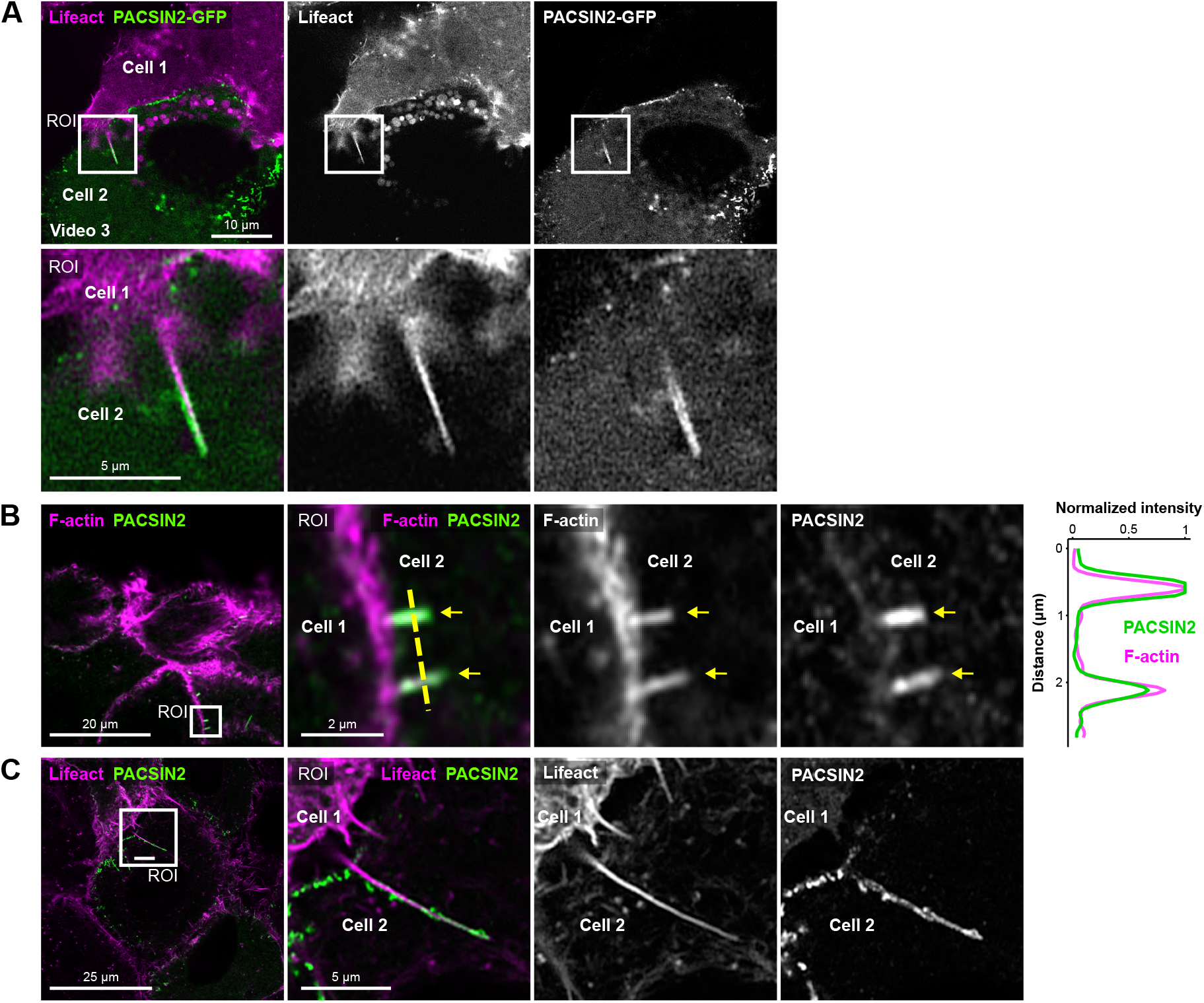
PACSIN2 is recruited to filopodia impact sites. (**A**) DCIS.com lifeact-RFP cells were co-cultured with DCIS.com parental cells transiently expressing PACSIN2-GFP. Live imaging was performed using Airyscan confocal microscopy. In this setup, PACSIN2-GFP-expressing cells are negative for lifeact-RFP, allowing unambiguous identification of the cellular origin of the PACSIN2 fingers. A representative field of view and a magnified region of interest (ROI) are shown. Scale bars: main image, 10 µm; ROI, 5 µm. (**B**) DCIS.com parental cells were cultured to confluence, fixed, and stained for F-actin and endogenous PACSIN2, then imaged with Airyscan confocal microscopy. A representative field of view and a magnified ROI are shown. Yellow dashed lines indicate the position of the line intensity profiles displayed adjacent to the microscopy images. Scale bars: main image, 20 µm; ROI, 2 µm. (**C**) DCIS.com lifeact-RFP cells were cultured to confluence, fixed, and stained for endogenous PACSIN2, followed by imaging with Airyscan confocal microscopy. A representative field of view and a magnified ROI are shown. Scale bars: main image, 25 µm; ROI, 5 µm. The raw images used to make this figure have been archived on Zenodo (Grobe et al., 2026).

**Supplementary Figure S6.**
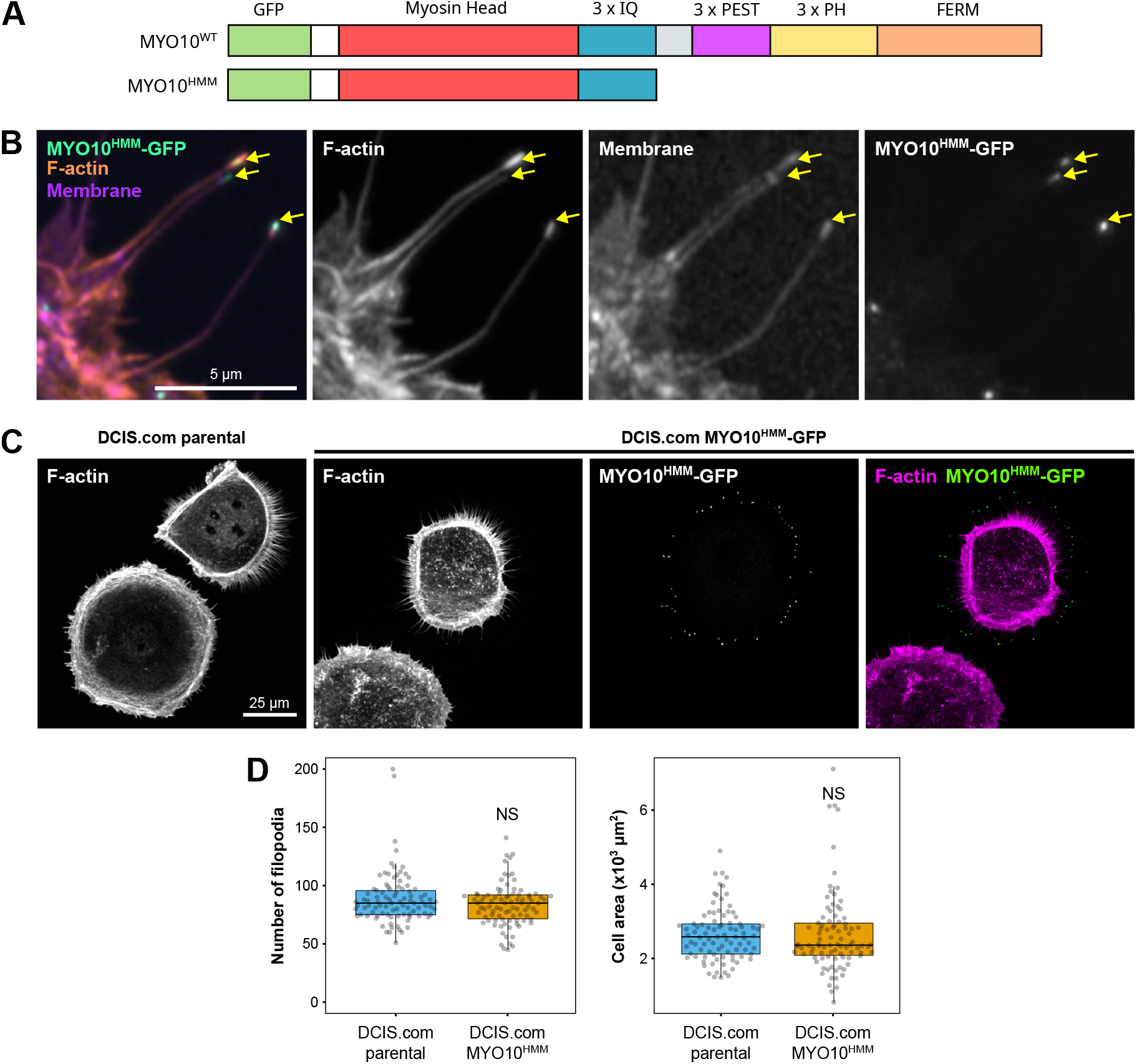
MYO10HMM-GFP localises to filopodia tips without inducing filopodia formation in DCIS.com cells. (**A**) Schematic illustration comparing the domain architecture of full-length wild-type MYO10 with the truncated MYO10HMM construct, which includes the head, neck, and coiled-coil domains but lacks the tail domain. (**B**) Representative Airyscan confocal image of DCIS.com cells expressing MYO10HMM-GFP, plated on fibronectin. Scale bar: 5 µm. (**C, D**) Parental and MYO10HMM-GFP DCIS.com cells were plated on fibronectin, fixed, and stained for F-actin. Samples were imaged using spinning disk confocal microscopy. (**C**) Representative images are displayed. Scale bar: 25 µm. (**D**) Quantification of filopodia number and cell spreading area in parental DCIS.com and MYO10HMM-GFP-expressing cells. (n = 90 cells from 3 independent experiments). Results are shown as boxplots, with whiskers extending from the 10th to the 90th percentiles. The boxes indicate the interquartile range, and a line within each box marks the median. Data points outside the whiskers are shown as individual dots. The raw images used to make this figure have been archived on Zenodo (Grobe et al., 2026).

**Supplementary Figure S7.**
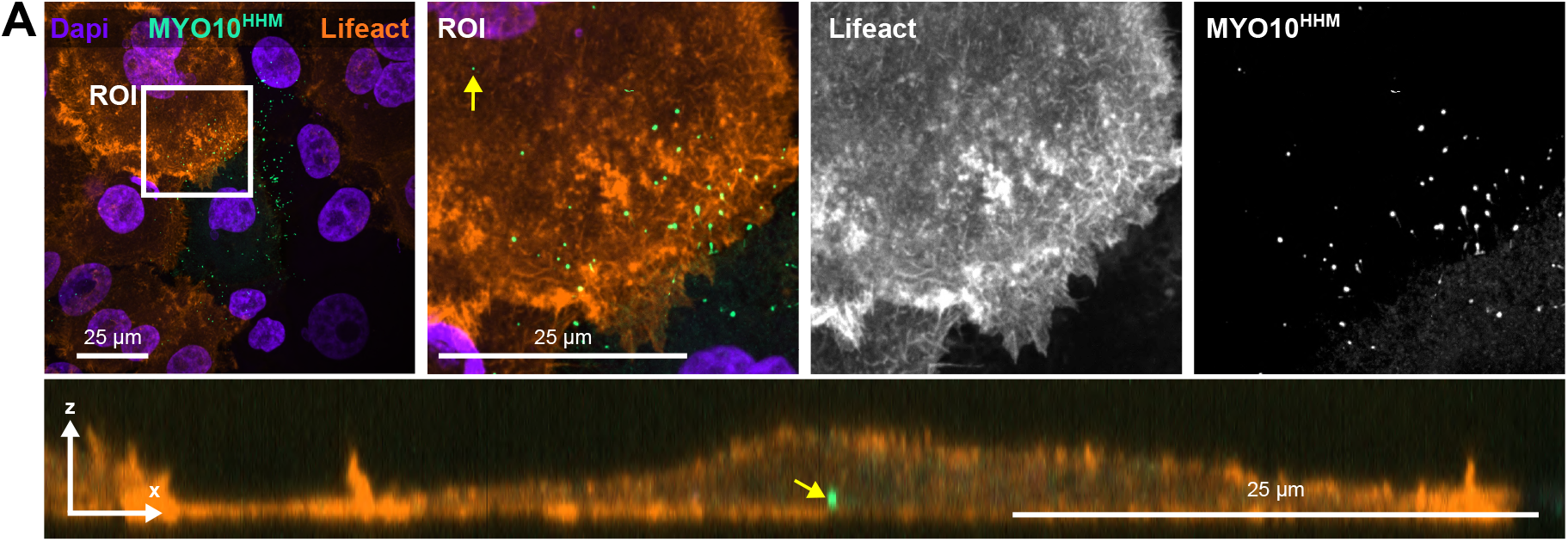
Intercellular filopodia tips can be internalised by neighbouring cells. (**A**) DCIS.com lifeact-RFP cells were co-cultured with DCIS.com MYO10HMM-GFP cells, fixed, stained using DAPI, and imaged using spinning disk confocal microscopy. Representative xy and xz planes are displayed. The yellow arrow highlights an internalised filopodia tip. Scale bars: 25 µm. The raw images used to make this figure have been archived on Zenodo (Grobe et al., 2026).

**Supplementary Figure S8.**
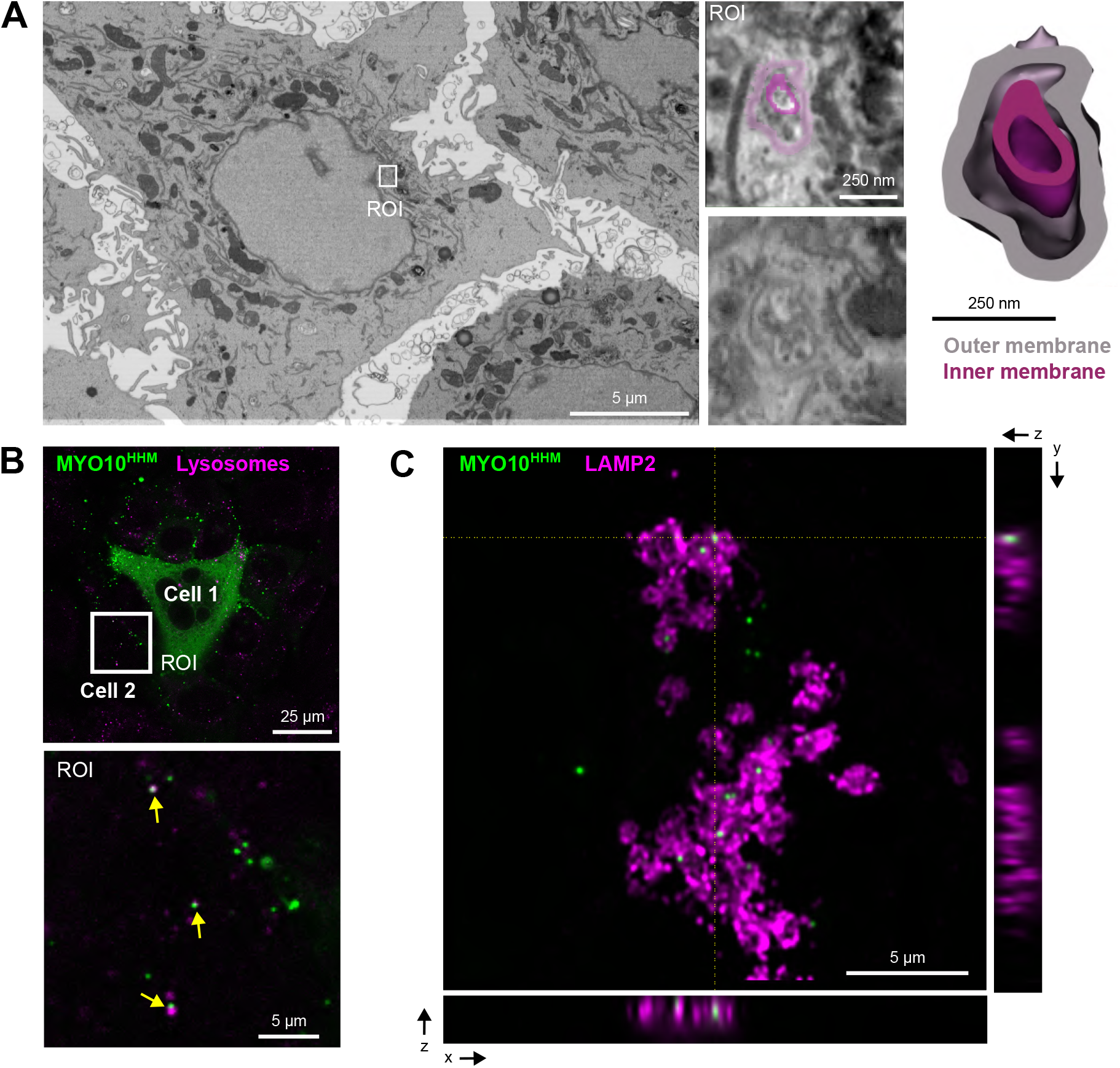
Internalised filopodia tips can fuse with lysosomes. (**A**) DCIS.COM cells expressing MYO10HMM–GFP were mixed with parental cells and cultured as a monolayer. After 24 hours, cells were fixed, labelled to visualise the plasma membrane and nuclei (DAPI), and imaged using high-resolution Airyscan confocal microscopy followed by focused ion beam–scanning electron microscopy (FIB–SEM). A MYO10HMM-GFP-positive punctum identified in the correlated EM volume is highlighted and displayed with its segmentation. Scale bars: 5 µm. (**B**) DCIS.com MYO10HMM-GFP and parental cells were mixed and imaged live after incubation with SiR-lysosome. A representative confocal image is shown. The white square indicates an ROI shown at higher magnification; yellow arrows highlight MYO10HMM–GFP-positive internalised filopodia tips overlapping with SiR-lysosome signal. Scale bars: main image, 25 µm; ROI, 5 µm. (**C**) DCIS.com MYO10HMM–GFP and parental cells were mixed and cultured in the presence of concanamycin A (125 nM, 24 hours). Cells were fixed, stained for LAMP2, and imaged using Airyscan confocal microscopy. Representative orthogonal views (xy, xz, and yz) are shown. Scale bar: 25 µm. The raw images used to generate this figure have been archived on Zenodo (Grobe et al., 2026).

